# Differences in *RAD51* transcriptional response and cell cycle dynamics reveal varying sensitivity to DNA damage among *Arabidopsis thaliana* root cell types

**DOI:** 10.1101/2024.02.15.580253

**Authors:** Konstantin Kutashev, Anis Meschichi, Svenja Reeck, Alejandro Fonseca, Kevin Sartori, Charles White, Adrien Sicard, Stefanie Rosa

**Affiliations:** Swedish University of Agricultural Sciences, Plant Biology Department, Uppsala, Sweden; Institute of Molecular Plant Biology, Department of Biology, Swiss Federal Institute of Technology Zürich, 8092 Zürich, Switzerland; Department of Cell and Developmental Biology, John Innes Centre, Norwich Research Park NR4 7UH, UK; Institut Génétique Reproduction et Développement (iGReD), Université Clermont Auvergne, UMR 6293, CNRS, U1103 INSERM, Clermont-Ferrand, France

**Keywords:** DNA damage, Double Strand Breaks, Homologous Recombination, *RAD51* transcription, Cell cycle arrest, Cell cycle checkpoints, *Arabidopsis thaliana*

## Abstract

- Throughout their lifecycle, plants are subjected to DNA damage from various sources, both environmental and endogenous. Investigating the mechanisms of the DNA damage response (DDR) is essential to unravel how plants adjust to the changing environment that can elicit varying amounts of DNA damage.
- Using a combination of state-of-the-art cell biology methods including whole-mount single-molecule RNA fluorescence *in situ* hybridization (WM-smFISH), allowing detection of individual mRNA molecules in intact plant tissue and plant cell cycle reporter lines we investigated how the transcriptional activation of a key homologous recombination (HR) gene, *RAD51*, occurs in response to increasing amounts of DNA damage in *Arabidopsis thaliana* roots.
- The results uncover consistent variations in *RAD51* transcriptional response and cell cycle arrest among distinct cell types and developmental zones. Furthermore, we demonstrate that DNA damage induced by genotoxic stress results in *RAD51* transcription throughout the whole cell cycle, dissociating its traditional link with S/G2 phases.
- This work advances the current comprehension of DNA damage response in plants showing quantitative differences in DDR activation. In addition, it reveals new associations with the cell cycle and cell types, providing crucial insights for further studies of the broader response mechanisms in plants.

## INTRODUCTION

Plants, due to their sessile nature, are constantly exposed to various DNA damaging agents from both the environment and endogenous processes. One of the most dangerous lesions that can occur on the DNA are double-stranded breaks (DSBs) (Vítor *et al*., 2020). The occurrence of this type of lesions requires immediate repair, resulting in DNA damage response (DDR) activation, recruitment of the DNA repair machinery to the lesion site and cell cycle arrest until the repair is complete (Preuss & Britt, 2003; Cools *et al*., 2011). In most cases, DSBs are repaired by one of two mechanisms: non-homologous end joining (NHEJ) or homologous recombination (HR) (West *et al*., 2004). HR is typically viewed as an error-free repair system that relies on an intact DNA strand acting as a template for reconstruction of the broken DNA strand (Schuermann *et al*., 2005).

Soon after the occurrence of DSB thousands of kilobases around the newly formed DSB are labeled by phosphorylated form of H2A.X histone variant (γ-H2AX) that participates in the early signaling of the lesion and recruitment of DNA repair machinery proteins (Rogakou *et al*., 1999; Stewart *et al*., 2003; Lang *et al*., 2012; Fan *et al*., 2022). Histone γ-H2AX levels were shown to correlate with DNA damage amounts (Friesner *et al*., 2005; Redon *et al*., 2009; Lee *et al*., 2019), and its dynamics of recruitment and loss are employed to measure DSB repair dynamics (Löbrich *et al*., 2010; Lee *et al*., 2019).

HR pathway is intricately connected to the S and G2 phases of the cell cycle, owing to its inherent need for an intact repair template, with the sister chromatid predominantly serving this function (Johnson, 2000; Saleh-Gohari, 2004; Saintigny *et al*., 2007; Goldfarb & Lichten, 2010; Bee *et al*., 2013). This regulation is accomplished through cell-cycle-linked transcriptional control of HR proteins and post-translational modifications of proteins during the S and G2 phases (Yata *et al*., 2012; Weimer *et al*., 2016; Lim *et al*., 2020). Cell cycle arrest is an integral part of the DDR, providing the necessary time for repair to take place thus ensuring the integrity of genetic material (Muschel *et al*., 1991; Raleigh & O’Connell, 2000; Chen *et al*., 2017). In *A. thaliana*, cell cycle arrest in response to DNA damage occurs mostly at G2/M and G1/S phase checkpoints (De Schutter *et al*., 2007; Cui *et al*., 2017; Cabral *et al*., 2020). Different sources of DNA damage arrest the cell cycle at different stages. For example, hydroxyurea or cadmium-induced DNA damage arrests the cell cycle at G1/S transition (Culligan *et al*., 2004; Cui *et al*., 2017), while γ-irradiation and aphidicolin arrest the cell cycle at G2/M transition (Culligan *et al*., 2004).

RAD51 protein plays a key role in repair via HR. RAD51 promotes essential strand-invasion step where resected 3’ single-stranded DNA end aligns with a homologous template, ensuring proper placement of broken DNA strand overhangs (Shinohara *et al*., 1992; Abe *et al*., 2005; Li *et al*., 2004; Su *et al*., 2017; Wang *et al*., 2014; Yu *et al*., 2023; Banerjee & Roy, 2021). The widespread presence of RAD51 homologs across various species underscores its fundamental functional significance (Bonilla *et al*., 2020). Mutation in *RAD51* gene is lethal in animals but dispensable for vegetative development in *A. thaliana* (Lim & Hasty, 1996; Tsuzuki *et al*., 1996; Li *et al*., 2004). Upon DNA damage, *RAD51* transcription is activated (Wang *et al*., 2014; Feng *et al*., 2017; Ryu *et al*., 2019; Da Ines *et al*., 2022) in a dose-dependent manner (Osakabe *et al*., 2005; De Schutter *et al*., 2007). RAD51 protein is subsequently loaded onto the lesion site by BRCA2 or CX3 complex (Wang *et al*., 2010; Su *et al*., 2017). This accumulation of RAD51 at the broken strand overhang (Flott *et al*., 2011; Biedermann *et al*., 2017; Da Ines *et al*., 2022) facilitates the search for a homologous donor template (Hicks *et al*., 2011; Coïc *et al*., 2011; Meschichi *et al*., 2022). The activity of RAD51 is tightly regulated at the post-translational level, serving as a substrate of multiple kinases in human (Sørensen *et al*., 2005; Chabot *et al*., 2019; Woo *et al*., 2021) and budding yeasts (Flott *et al*., 2011; Woo *et al*., 2020, 2021). Although RAD51 phosphorylation by the cyclin kinase CDKB1-CYCB1 complex was reported *in vitro* for *A. thaliana*, its exact function remains elusive. Nevertheless, it is highly likely that this process is linked to RAD51 activation and recruitment to the DNA double-strand break sites, as evidenced by compromised RAD51 foci formation in *cycb1;1* mutants (Weimer *et al*., 2016).

Most plant studies have analyzed *RAD51* expression using bulk measurements, combining material from multiple plants (Wang *et al*., 2014; Ryu *et al*., 2019). Although this approach is suitable for many purposes, it does not allow determining how gene expression is tuned at the level of individual plants, tissues, or cell types. In this study, we used single-molecule RNA FISH (smFISH) (Duncan *et al*., 2017; Zhao *et al*., 2023) to quantify the transcriptional response of *RAD51* at the cellular level to increasing amounts of DNA damage induced by DNA damaging agent zeocin (Adachi *et al*., 2011). Our findings show a positive correlation between *RAD51* transcription and increasing amounts of damage, and we demonstrate that *RAD51* mRNA output reaches a maximum at the cellular level upon surpassing a certain damage threshold. Notably, *RAD51* transcriptional response was different between root cell types and developmental zones. Our data also shows prominent *RAD51* transcription outside S/G2 cell cycle phases under DNA damage, challenging the proposed strict association between HR and S/G2 phases.

## MATERIALS AND METHODS

### Plant material

All *Arabidopsis thaliana* lines used in this study were derived from Columbia (Col-0) ecotype. Transgenic lines used in this study come from the following sources: RAD51-GFP line (Da Ines *et al*., 2013), Cytrap (Aki & Umeda, 2016), PlaCCI and CDT1-CFP lines (Desvoyes *et al*., 2020).

### Plant growth

Arabidopsis seeds were surface sterilized in 5% (v/v) sodium hypochlorite for 5 min and rinsed three times in sterile distilled water. Seeds were then stratified for 2 days at 4°C before germination in a growth chamber in a vertically oriented Petri dish containing 1% plant agar (Duchefa Biochemie, P1001.1000) MS medium plate, pH 5.7 (Gamborg *et al*., 1976). Plants were grown under a photoperiod of 16 hours day and 8 hours night and a temperature cycle of 22°C during the day and 20°C during the night.

### Expression analysis using real-time RT-PCR (qPCR)

For total *RAD51* transcript analysis, 10-day-old *A. thaliana* (Col-0) seedlings were measured by qPCR. 9-day-old seedlings were transferred onto 1% plant agar MS medium plates containing different concentrations of zeocin (0 μM, 10 μM, 50 μM, 170 μM) (Gibco, 10072492). Seedling roots were cut with a razor blade and collected after overnight zeocin exposure. A total of 0.1g of roots per zeocin concentration was used. RNA was isolated using Quiagen RNeasy Plant Mini kit (Quiagen, 74904). RNA concentration was measured using Nanodrop ND-1000 spectrophotometer. A total of 1 μg of RNA was treated with DNase (Thermo Fisher, EN0521) and reverse transcribed with Reverse Transcriptase (Thermo Fisher, EP0441). This template was then used to quantify relative mRNA abundance using the SensiMix SYBR Low-ROX kit (Bioline), a LightCycler® 480 (Roche) and the primers described below. *RAD51* expression was analyzed using normalization to *PP2A* gene using following primers: *Rad51* forward GCGCAAGTAGATGGTTCAGC, *Rad51* reverse TTCCTCAACGCCAACCTTGT, *PP2A* forward TAACGTGGCCAAAATGATGC, *PP2A* reverse GTTCTCCACAACCGCTTGGT. Reactions were performed in triplicate, results were calculated using the 2^-ΔΔ*CT*^ method, standard deviation values shown on a graph.

### Single molecule fluorescence *in situ* hybridization (smFISH) on root squashes

smFISH was performed on 5-6 days old seedlings according to previously published protocol (Duncan *et al*., 2017) using probes designed against *RAD51* and *PP2A* genes (listed in Supplementary Table 2). Seedlings were transferred onto MS medium plates containing selected concentrations of zeocin (Gibco, 10072492) or no zeocin for control sample. Seedlings were collected after overnight zeocin exposure and treated further according to protocol.

### Immunodetection

5-6 days old seedlings were transferred onto MS medium containing selected concentrations of zeocin overnight. Roots were then cut off from seedlings using a razor blade and fixed in 4% paraformaldehyde solution for 30 minutes in glass dishes. Roots were then washed twice with 1x PBS. 5 roots were then arranged on a slide in similar orientation, covered by a glass coverslip and squashed manually by applying pressure on coverslip. The slide was then submerged in liquid nitrogen until freezing and taken out, coverslip was then removed using a razor blade. Slides were left to dry at room temperature for 30 minutes. Samples were rinsed with 1x PBS three times and incubated with blocking buffer (0.5% BSA (Sigma-Aldrich A7030) in 1x PBS) in humid chamber at 37°C for 30 minutes. To ensure minimal disturbance of the sample we used small pieces of polypropylene waste bags instead of glass coverslips at all incubation stages of the protocol. Excess blocking buffer was removed, samples were incubated at 37°C overnight in a humid chamber with ɣH2AX primary antibody (Charbonnel *et al*., 2010), provided by Charles White. Antibody was diluted 1:700 in 0.5% BSA. Slides were then rinsed with PBST buffer three times (1x PBS, 0,01% Tween20 (Sigma-Aldrich 8.22184)) and incubated with PBST buffer for 5 min. Secondary antibody (Agrisera, AS09633) diluted 1:200 in 0.5% BSA was then applied, and samples were incubated in a humid chamber at 37°C for 2 hours. Slides were rinsed three times with PBST buffer and incubated with 1x PBS buffer 2x 5 min. Excess buffer was removed, and samples were mounted in Vectashield medium (Vector laboratories, H-1000) containing DAPI diluted 1:1000 (Thermo Fisher, 62248).

### Sequential smFISH and immunodetection

SmFISH and immunodetection protocols were performed sequentially in the described order. SmFISH in root squashes was performed first according to the protocol mentioned above. After imaging the coverslips were gently removed using additional volumes of 1x PBS. Samples were rinsed with 1x PBS three times and samples were processed according to immunodetection protocol above.

### Whole-mount smFISH (WM-smFISH)

WM-smFISH was performed on 5-6 days old seedlings according to previously published protocol (Zhao *et al*., 2023) using probes designed against *Rad51* gene (listed in Supplementary Table 1). Seedlings were transferred onto MS medium plates containing selected concentrations of zeocin (Gibco, 10072492) or no zeocin for control sample. Seedlings were collected after overnight zeocin exposure and treated further according to protocol.

### Sequential WM-smFISH and 5-ethynyl-2’-deoxyuridine (EdU) labeling

5-6 days old seedling were first transferred onto the MS medium containing zeocin for 10 hours. Seedlings were then transferred onto MS medium containing same concentration of zeocin and 20 μM EdU (Invitrogen, A10044) for two hours. WM-smFISH was performed first according to the described protocol. After imaging, coverslips were gently removed from the samples using additional volumes of 2x SSC buffer. Samples were then rinsed with 2x SSC buffer three times and incubated with 3% BSA in 1x PBS solution at 37°C in a humid chamber for 15 minutes. Samples were incubated with Click-iT reaction cocktail (Invitrogen C10269) mixed according to the manufacturer’s instructions with addition of Alexa Fluor 488 azide (Thermo Fisher, A10266), 500x dilution. Samples were then rinsed and incubated with a wash buffer (10% formamide (Thermo Scientific, 17899) and 2xSSC) for 5 minutes. Samples were incubated with SCRI Renaissance 2200 solution (Musielak *et al*., 2015) for 15 minutes at 37°C in a humid chamber. Slides were rinsed and incubated for 5 min in the wash buffer. Samples were then mounted in a drop of Vectashield medium.

### *RAD51* mRNA half-life quantification

5-6 days old seedling were transferred onto MS medium containing 10μM zeocin for selected time periods: 12, 10, 8, 6 hours. Seedlings exposed to zeocin for 10, 8 and 6 hours were then transferred to MS medium containing zeocin with Actinomycin D (Thermo Fisher, J60148.LB0) or zeocin with DMSO (Sigma-Aldrich, D4540) for 2, 4 and 6 hours accordingly. Seedlings were then collected and processed according to smFISH protocol for root squashes using probes for *RAD51* gene. The decay rate (kdecay) for *RAD51* and then its half-life (t1/2) were calculated by adjusting the number of molecules per cell (N) counted in the smFISH images as an exponential function of time (t). The mathematical adjustment for N(t) was developed in R assuming a constant decay rate, according to the function: N(t) = e^-kdecay^ * ^t^, then the half-life was calculated using the formula: ln(2)/kdecay) (Narsai *et al*., 2007; Sorenson *et al*., 2018).

### Image acquisition

Samples were imaged on Zeiss LSM780 and LSM800 inverted confocal microscopes (Zen Black Software) using a 63X water-immersion objective (1.20 NA). smFISH on root squashes imaging was performed using widefield mode, we used a cooled quad-port CCD (charge-coupled device) ZEISS Axiocam 503 mono camera. A series of optical sections with z-steps of 0.22 μm were collected throughout the whole cell volume. For DAPI imaging an excitation filter of 335-383 nm was used and emission was collected at 420 - 470 nm. Quasar570 fluorescent probes were imaged using 533-558 nm excitation filter and 570-640 nm signal detection range. For immunostaining experiments were did not use widefield mode, for DAPI signals excitation line of 405 nm was used with emission detection at 410-600 nm. For GFP signals of labelled histone ɣH2AX excitation line of 488 nm and emission at 490 - 540 nm settings were used. Imaging was performed in a manually adjusted single plain selected to have a maximal number of nuclei in focus.

WM smFISH imaging was performed in confocal mode using a 63X water-immersion objective (1.20 NA). For SCRI Renaissance 2200 imaging we used a 405 nm laser line and and emission was collected at 410-600 nm. Quasar570 probe signals were captured with 561 nm excitation line and emission collection at 565-700 nm. CFP signals were imaged using 455 nm excitation line and emission detection at 460-600 nm.

### Image analysis

#### smFISH

Nuclei and cellular outlines in smFISH were defined using CellProfiler software (Stirling *et al*., 2021). RNA foci were detected and counted using FISH-quant-v3 (Mueller *et al*., 2013) in Matlab. First, the “cell segmentation” tool was used to generate text files with the outline coordinates for the nuclei and cell masks. The outlines were uploaded, and images were pre-processed for increasing their signal-to-noise ratio though a dual-Gaussian filtering followed by a Gaussian Kernel. Dots were detected in the filtered image, first pre-detecting fluorescent foci with fluorescence over a threshold. Then, the pre-detected dots were fitted to a Gaussian fluorescence based on a point-spread function. Images were analyzed in the batch mode, and false positives were removed in the end by thresholding the Sigma-XY, amplitude, and pixel-intensity parameters to Gaussian distributions.

#### WM-smFISH

Cell segmentation was performed with Cellpose software (Stringer *et al*., 2021), using an algorithm trained by us. RNA foci were detected and counted using FISH-quant-v3 (Mueller *et al*., 2013) as described above. For RAD51-GFP line the signal intensities of both mRNA and protein channels were quantified in CellProfiler software (Stirling *et al*., 2021). Colocalization analysis and heatmap visualization was performed using CellProfiler software (Stirling *et al*., 2021).

Images from EdU staining, Cytrap and PlaCCI lines were analysed manually using ImageJ software.

#### Correlation analysis of ɣH2AX signal and RAD51 transcription

Data on *RAD51* transcription and H2AX levels were collected from the same cells for correlation analysis. ɣH2AX integrated density was measured using ImageJ software and normalized to DAPI integrated density. The number of detected *RAD51* mRNA molecules was normalized by the cell area to correct for cell size difference. Values obtained for both parameters were log transformed. Data was visualized and correlation was evaluated using R studio ggplot2 package.

## RESULTS

### RAD51 transcriptional response to increasing DNA damage levels

To elucidate *RAD51* transcriptional response to DNA damage we first assessed *RAD51* mRNA levels on roots from Col-0 plants treated with increasing concentrations of the DSB-inducing agent zeocin (0 μM, 10 μM, 50 μM, 170 μM) using real-time quantitative PCR (RT-qPCR). The results demonstrated an increase in *RAD51* mRNA quantities with increasing zeocin concentrations (**Fig. 1a**). To investigate *RAD51* transcriptional upregulation as a function of DNA damage at the cellular and tissue level we employed smFISH (Duncan *et al*., 2016), a method that allows absolute quantification of transcripts in individual cells (**Fig. 1b, c(i)**), using the same material preparation as for immunodetection. Consistent with the qPCR data (**Fig. 1a**) smFISH results revealed an increase in the total number of *RAD51* mRNAs in root tissue with increasing zeocin concentrations (**Fig. 1d**). Of note, the total number of *RAD51* mRNAs did not seem to increase in direct proportion to the concentration of zeocin. To evaluate the increase in DNA damage levels corresponding to increasing zeocin concentrations, we quantified γ-H2AX levels by immunodetection as a proxy marker for DSB levels in individual root cells. Single cell spreading achieved by root squashing facilitated antibody penetration required for immunodetection (**Fig. 1c(ii)**). The results showed an accumulation of γ-H2AX in the nuclei of root cells in response to growing zeocin concentrations (**Fig. 1e**), confirming the increase in DSBs. The observed increase in γ-H2AX accumulation was not directly proportional to the increase in zeocin concentration as the number of *RAD51* mRNAs. To assess the direct relationship between *RAD51* transcription and the extent of DNA damage within individual cell, we performed a sequential *RAD51-*smFISH/ɣH2AX-immunodetection protocol on cells obtained from root squashes and evaluated mRNA and DNA damage levels on the same cells (**Fig. 1c**). This analysis revealed a positive correlation between the number of *RAD51* mRNA molecules per cell and the ɣ-H2AX levels with a correlation coefficient R=0.62 (p<1.4e^-14^) (**Fig. 1f**). Our analysis indicated that the interaction between the two variables is best described by a linear model with deviance of fit (DOF) value of 18.44615. DOF value of the exponential model, indicating a potential limit to the possible number of *RAD51* mRNAs per cell, was however only slightly higher, 18.99684 (**Fig. S1**). Importantly, the mRNA counts for the house-keeping gene *PP2A* remained constant across zeocin concentrations (**Fig. S1**), confirming the specific *RAD51* upregulation with increasing damage.

**Fig. 1.**
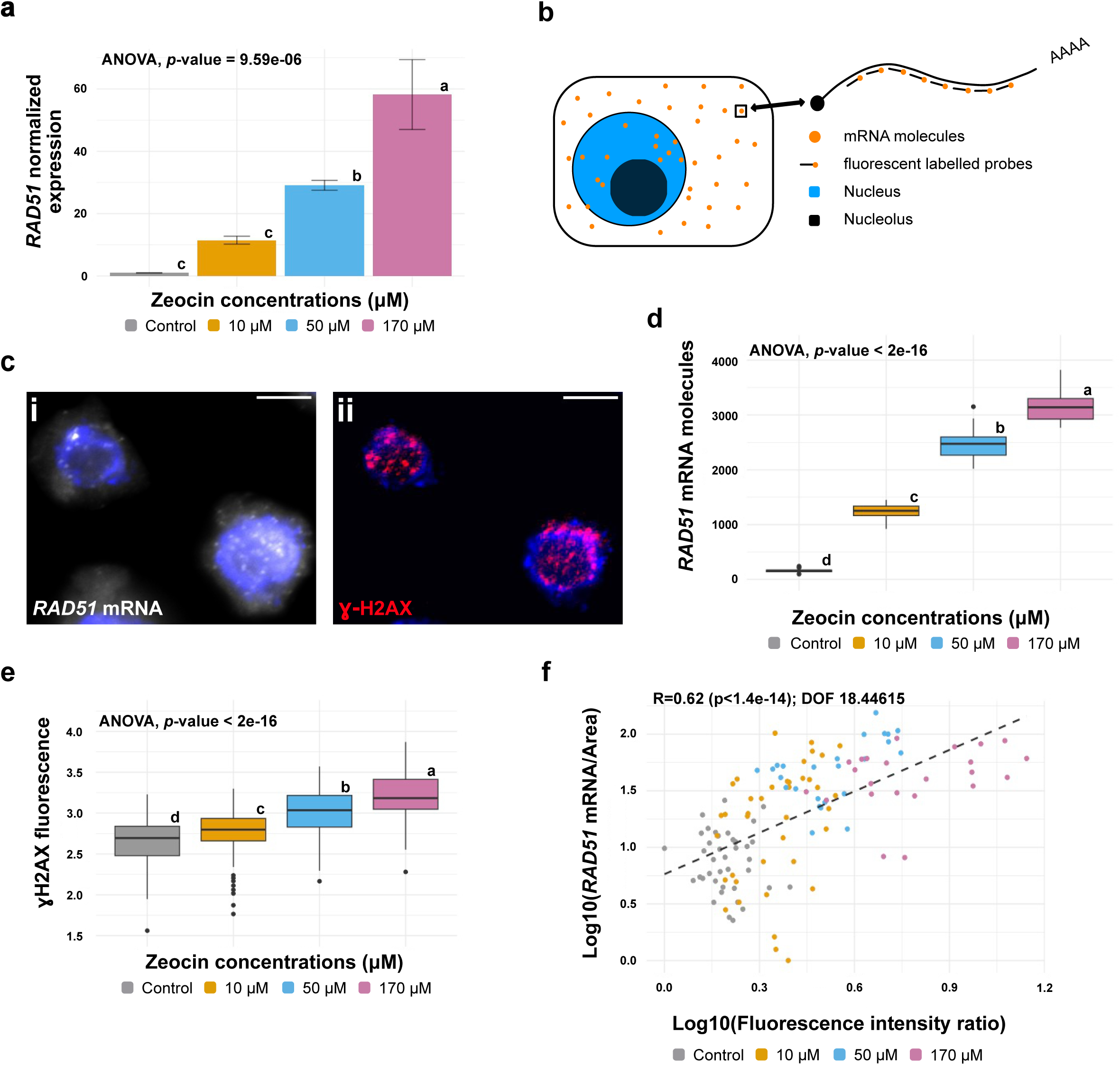
*RAD51* transcription and ɣ-H2AX accumulation in response to increasing DNA Damage in *Arabidopsis thaliana* root squashes. (**a**) qPCR quantification of *RAD51* transcriptional response in roots after exposure to 0 μM, 10 μM, 50 μM, 170 μM concentrations of zeocin. *RAD51* expression measured relative to *PP2A* gene as a control. ANOVA revealed statistically significant difference in *RAD51* expression by zeocin concentration (F(3)=78.05, *p*=9.59e-06)). Letters indicate results of TukeyHSD test with 95% confidence level. Error bars indicate standard deviation. (**b**) Schematic image of single mRNA molecule labelling using multiple fluorescent probes by smFISH protocol. (**c**) Images acquired using sequential smFISH and immunodetection protocol in squashed root cells after 50 μM zeocin exposure. (i) *RAD51* mRNA detection via smFISH, nuclei counterstained with DAPI (blue). (ii) ɣ-H2AX immunodetection (red) and DAPI (blue) signals. Scale bars, 5 μm. (**d**) Total number of *RAD51* mRNA molecules detected in 30 randomly selected cells after zeocin exposure from dataset containing (n=160, n=116, n=60, n=96 cells for 0 μM, 10 μM, 50 μM, 170 μM zeocin respectively), subset was selected 30 times. ANOVA revealed statistically significant difference in *RAD51* mRNA molecule number by zeocin concentration (F(3)=1497, *p*<2e-16)). Letters indicate results of TukeyHSD test with 95% confidence level. (**e**) ɣ-H2AX fluorescence signal intensities measured after exposure to 0 μM, 10 μM, 50 μM, 170 μM concentrations of zeocin in squashed roots. Values represent nuclear signal fluorescence intensity measured as lg (Integrated Density). ANOVA revealed statistically significant difference in ɣ-H2AX fluorescence by zeocin concentration (F(3)=175.4, *p*<2e-08)) in our measurements (n=236, n=145, n=226, n=182 cells for 0 μM, 10 μM, 50 μM, 170 μM zeocin respectively). Letters indicate results of TukeyHSD test with 95% confidence level. (**f**) Correlation analysis between the number of *RAD51* mRNA molecules and ɣ-H2AX signal intensity in individual cells of squashed roots with linear model fit. Number of *RAD51* transcripts normalized by corresponding cell area, log10 of this value used for the corresponding axis. ɣ-H2AX fluorescence intensity measured as lg(Integrated Density) with prior normalization to DAPI Integrated density. Correlation coefficient (R) and p-value shown on a graph. DOF indicates deviance of fit calculated for the model. Dataset contains n=40, n=40, n=25, n=22 measurements for 0 μM, 10 μM, 50 μM, 170 μM zeocin respectively.

### Cell-to-cell variability in RAD51 transcriptional response

To unravel potential differences in *RAD51* transcriptional activation between different cell types and developmental zones of the root we performed recently developed whole-mount smFISH (WM-smFISH) protocol (Zhao *et al*., 2023). This method overcomes the limitations of traditional root squash sample preparation, enabling the assessment of transcript numbers within intact tissues (**Fig. 2a, Fig. S2**). Heatmaps of *RAD51* mRNA molecules per cell were generated to visualize number of transcripts per cell across root tissues (**Fig. 2b**). The results revealed that the number of transcribing cells as well as the number of *RAD51* mRNA molecules detected per cell increases in response to increasing zeocin concentrations. This pattern is further evident in the histogram quantification (**Fig. 2c**), depicting a progressive rise in the number of actively transcribing cells with increasing zeocin concentrations. This data also indicates a possible upper boundary on the number of *RAD51* mRNA molecules per cell. Indeed, one cannot observe a large difference in mRNA numbers per cell between 50 μM and 170 μM zeocin concentrations despite the large increase in concentration, as visualized on heatmaps (**Fig. 2b**) and in a graph form (**Fig. 2c, d**).

**Fig. 2.**
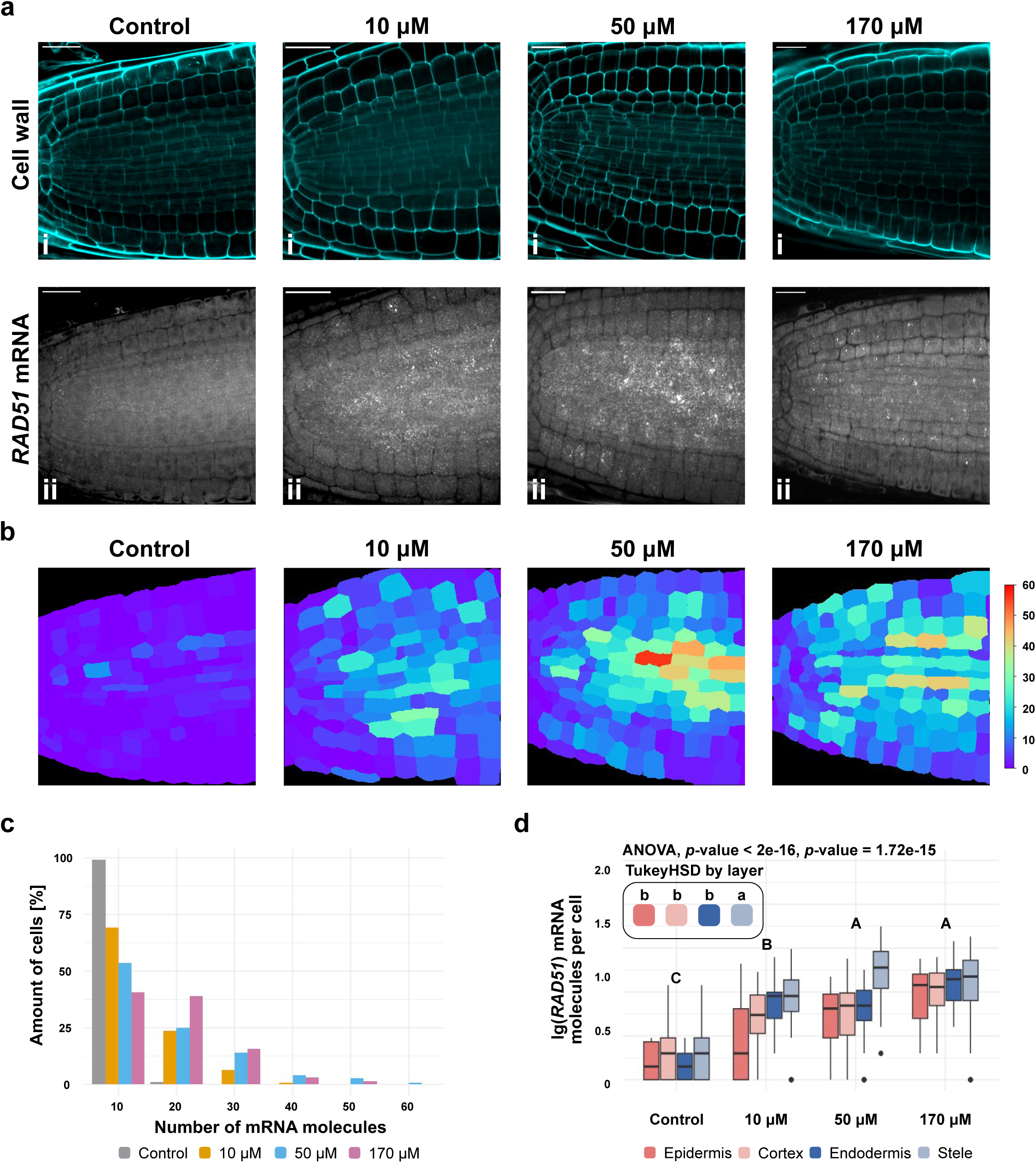
RAD51 mRNA transcriptional response in different cell types of *Arabidopsis thaliana* root. (**a**) Representative images of whole-mount smFISH for *RAD51* mRNA in Col-0 roots after exposure to 0 μM, 10 μM, 50 μM, 170 μM concentrations of zeocin. (i) Images of cell wall staining using Renaissance 2200 dye. (ii) Images of *RAD51* mRNA detection. Scale bars, 20 μm. (**b**) Heatmaps representing quantification of *RAD51* mRNA molecules detected in individual cells. (**c**) Frequency distribution of *RAD51* mRNA molecules per cell, for plants treated with different zeocin concentrations. Bin groups created using a step of 10 transcripts. (**d**) Number of *RAD51* mRNA molecules per cell in each of the selected cell types (Epidermis, Cortex, Endodermis, Stele) after exposure to 0 μM, 10 μM, 50 μM, 170 μM concentrations of zeocin. Two-way ANOVA revealed statistically significant difference in *RAD51* molecule number by both zeocin concentration (F(3)=119.93, *p*=2e-16)) and cell lineage (F(3)=25.39, *p*=1.71e-15)). Letters indicate results of TukeyHSD test of two-way ANOVA results with 95% confidence level. Measurements for (**c**) and (**d**) performed using data from images (**a**) and (**b**).

Importantly, our results indicate substantial variability among root cells in their sensitivity to DSBs induced by zeocin, as revealed by the non-uniform heatmaps. Some cells exhibited a strong transcriptional response even at a 10 μM zeocin concentration, with mRNA counts comparable to those induced by 50 μM and 170 μM (**Fig. 2b, c**). Conversely, certain cells displayed low mRNA counts even after exposure to 50 μM and 170 μM zeocin (**Fig. 2b, c**). To discern potential distinctions between cell types, we plotted the number of *RAD51* mRNAs in different root cell types (Epidermis, Cortex, Endodermis, and Stele). The results demonstrated that *RAD51* transcriptional response within stele cells was distinct from the other root cell types analyzed showing higher per cell mRNA output (**Fig. 2d**). The comparison also indicated no difference in mRNA counts per cell between 50 μM and 170 μM zeocin treated samples (**Fig. 2d**), potentially arguing in preference of previously proposed limit to per cell transcript output (**Fig. S1a**). Notably, across developmental regions *RAD51* transcriptional output decreased in the elongation zone in comparison with the meristem region (**Fig. S3**) consistent with previous reports (Da Ines *et al*., 2013).

### Quantification of RAD51 protein levels per cell

Next, we aimed to investigate the relationship between *RAD51* mRNA and protein levels per cell to assess the extent to which the rise in mRNA numbers aligns with the resultant protein quantity. For that we performed WM-smFISH on RAD51-GFP line (Da Ines *et al*., 2013) and quantified mRNA and protein mean fluorescence intensity per cell as described previously (**Fig. 3a**) (Zhao *et al*., 2023). Similarly to Col-0 plants, *RAD51* mRNA levels per cell exhibited an increase with increasing zeocin concentration in RAD51-GFP line (**Fig. 3a(ii, iv); Fig. 3b**). The results argue again in favor of linear growth of *RAD51* transcriptional response. However, we still observed that the increase in fluorescence was not isometric to the growth in zeocin concentration and substantial number of values between 50 μM and 170 μM zeocin treated samples were overlapping (**Fig. 3b**). The growth trend was similar for RAD51-GFP protein (**Fig. 3a(iii, v); Fig. 3c**). Of note, measured fluorescence signal intensity did not increase further after 50 μM zeocin concentration, suggesting a limit to RAD51 protein amount that can be present in the cell (**Fig. 3c**). Intriguingly, heatmaps evaluating ratio between mRNA and protein levels revealed slight differences between cells in terms of mRNA and protein accumulation (**Fig. 3a(vi); Fig. S4-6**). In line with our previous observations, mRNA molecules seem to have a higher abundance in stele (**Fig. 3a(ii, iv), a(vi); Fig. S4-6**). RAD51-GFP protein accumulation, on the other hand, was more prevalent in the cortex and epidermis of the root tip (**Fig. 3a(iii, v), a(vi); Fig. S3-5**). This differential accumulation between mRNA and protein among different cell types is intriguing and could suggest protein movement or differential degradation between cells but more investigation to validate these hypotheses would be required.

**Fig. 3.**
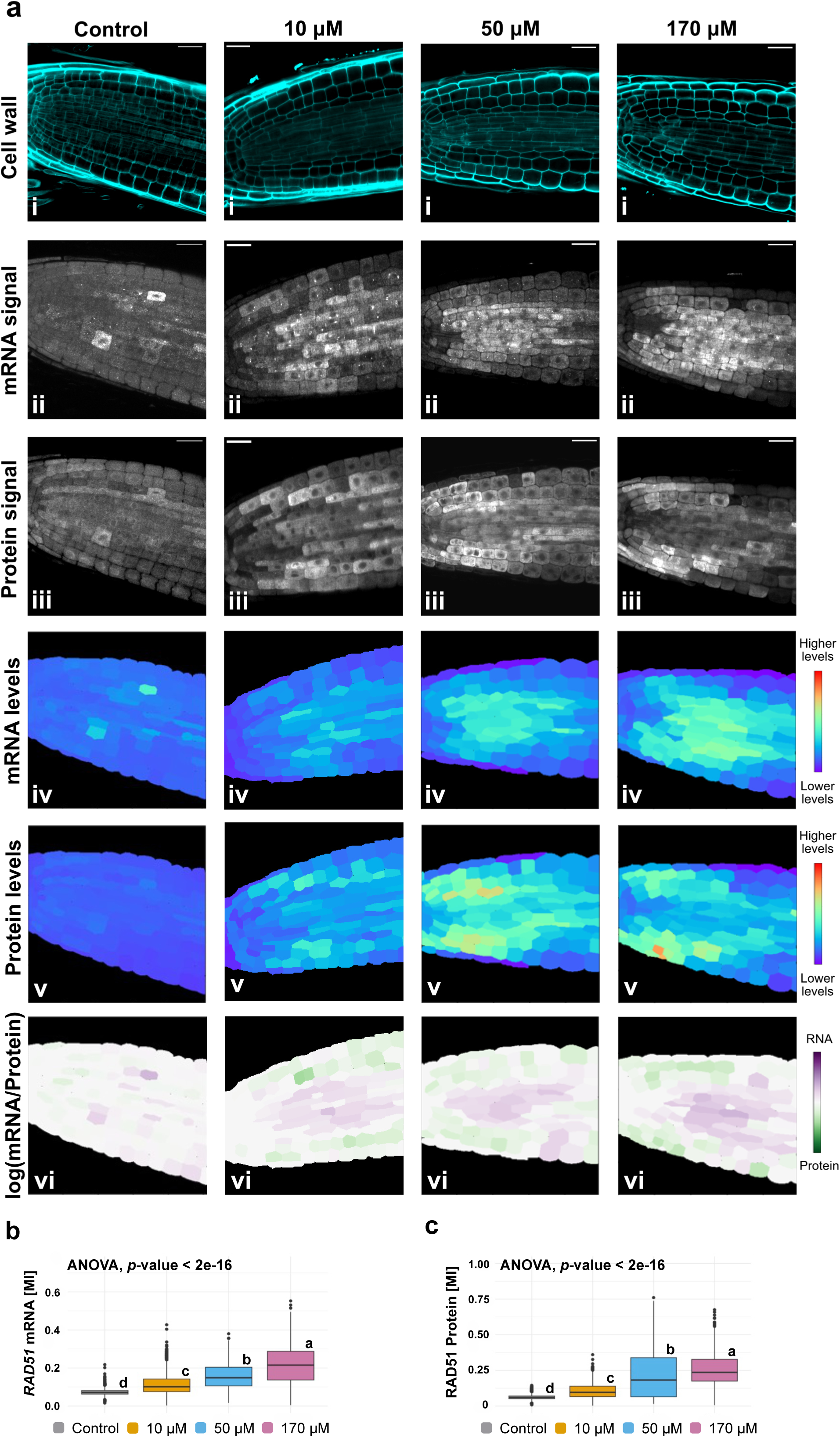
Simultaneous detection and quantification of *RAD51* mRNA and protein in response to increasing DNA damage in *Arabidopsis thaliana* RAD51-GFP line roots. (a) Representative confocal images and quantification of *RAD51* mRNA (ii) and RAD51-GFP protein signals (iii) after exposure to 0 μM, 10 μM, 50 μM, 170 μM concentrations of zeocin. (i) Imaging of cell wall staining using Renaissance 2200 dye. (ii, iii) Imaging of (ii) RAD51 mRNA by smFISH and (iii) RAD51-GFP signals. (iv, v) Heatmaps representing the levels of the corresponding mean signal intensity per cell (MI). (iv) Signal intensity of *RAD51* mRNA molecules. (v) Signal intensity of RAD51-GFP protein. (vi) Heatmaps representing the ratio between the *RAD51* mRNA and RAD51-GFP protein signal intensities in each cell. Scale bars, 20 μm. (b) *RAD51* mRNAs per cell after exposure to 0 μM, 10 μM, 50 μM, 170 μM zeocin. ANOVA revealed statistically significant difference in *RAD51* mRNA signal mean intensity by zeocin concentration (F(3)=558.7, p<2e-16)). Letters indicate results of TukeyHSD test with 95% confidence level. (c) RAD51-GFP mean signal intensity per cell after exposure to 0 μM, 10 μM, 50 μM, 170 μM zeocin. ANOVA revealed statistically significant difference in RAD51-GFP signal mean intensity by zeocin concentration (F(3)=547.2, p<2e-16)). Letters indicate results of TukeyHSD test with 95% confidence level. Graphs on (b) and (c) created using dataset from several images containing (n=640, n=1391, n=854, n=457 cells for 0 μM, 10 μM, 50 μM, 170 μM zeocin respectively) individual cell measurements.

### RAD51 transcription through the cell cycle

*RAD51* transcription is typically linked to the S and G2 phases of the cell cycle, motivated by the requirement of homologous DNA sequences during repair through HR (Schuermann *et al*., 2005; Goldfarb & Lichten, 2010). Given the very high proportion of cells with *RAD51* mRNAs signals in the zeocin-treated samples, we expected a considerable number of cells arrested at the S or G2/M checkpoints (Osakabe *et al*., 2005; De Schutter *et al*., 2007). To evaluate the cell cycle arrest in roots treated with zeocin, we conducted EdU staining to label cells that went through S-phase in a sequential smFISH/EdU protocol (**Fig. 4a-c**). Our results revealed a drastic decline in the number of EdU-positive cells with increasing zeocin concentration, with almost no labeled cells at 50 μM and 170 μM concentrations, indicating a strong cell cycle arrest in these samples (**Fig. 4b, 4d, Fig. S7**). EdU-positive cells tend to be most abundant in the root stele, possibly explaining higher *RAD51* transcript output in this part of the root as it is usually associated with S/G2 phases of the cell cycle **(Fig. S7a, S7b, S7c).** Moreover, two-way ANOVA revealed that the observed variations in EdU-positive cell numbers can be explained by both zeocin concentration and the cell type with significant interaction between the two parameters (*p* = 4.94e^-15^) (**Fig. S7b**). Subsequent pairwise comparisons revealed statistically significant changes in cell numbers between the concentrations only in the root stele. Importantly, comparing EdU labelling with *RAD51* smFISH signals revealed EdU stained cells with modest *RAD51* mRNA presence next to cells with no EdU signal and abundant number of *RAD51* mRNA molecules (**Fig. 4c**). This observation potentially challenges the strict dependency of *RAD51* transcription on the S/G2-phase of the cells. Of note, EdU signals were observed in the elongation zone of the root at 50 μM and 170 μM concentrations of zeocin, revealing distinct responses across the various root developmental zones (**Fig. S7c**).

**Fig. 4.**
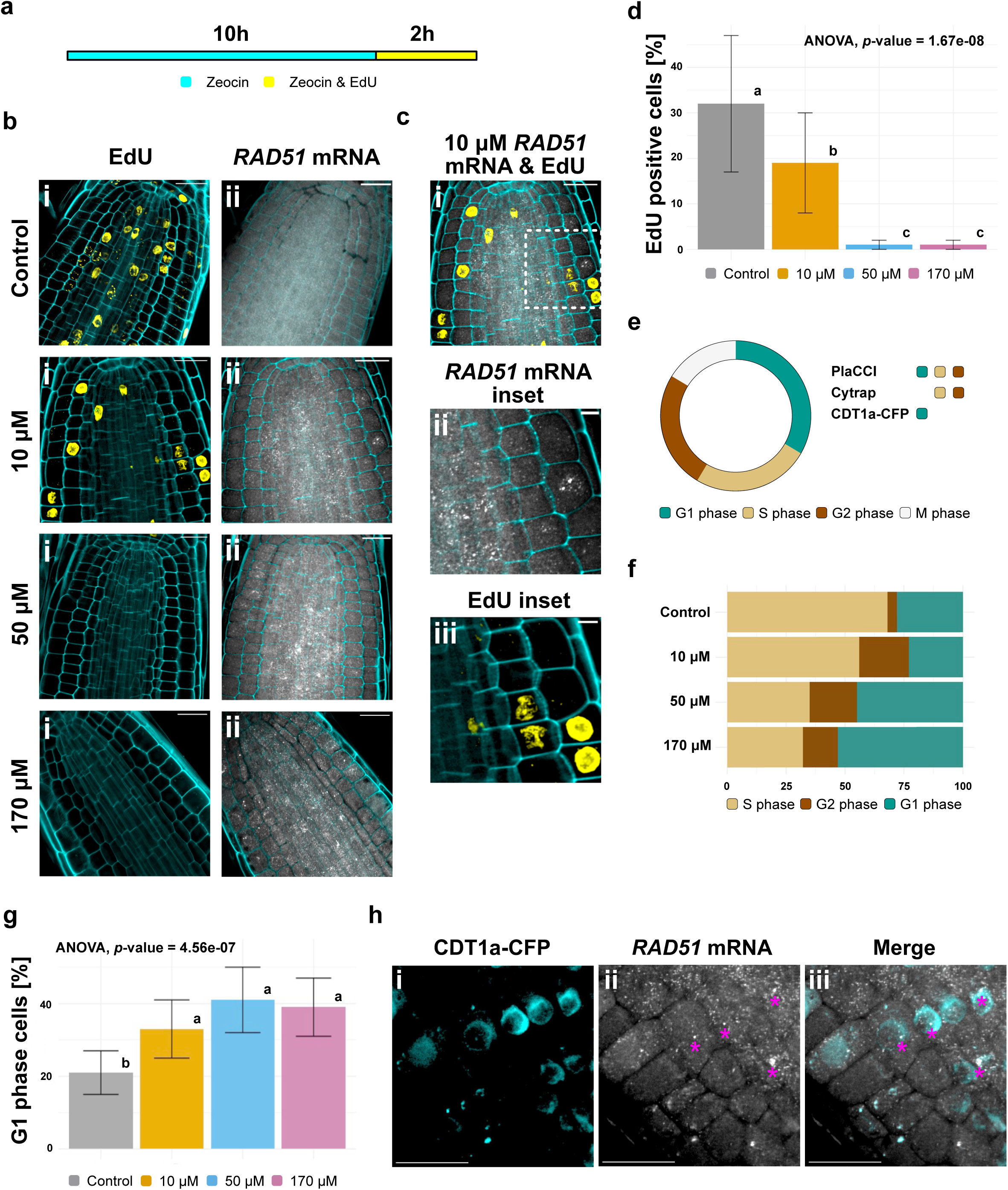
Dynamics of *RAD51* transcription throughout the *Arabidopsis thaliana* cell cycle. (**a**) Scheme of experimental setup used for quantification of S-phase cells. Seedlings were treated with different concentrations of zeocin concentrations for 10 hours, followed by additional treatment with zeocin and EdU for two hours. (**b**) Confocal images of roots acquired using sequential WM-smFISH/EdU staining protocol after exposure to 0 μM, 10 μM, 50 μM, 170 μM zeocin. Cell wall staining using Renaissance 2200 dye. (i) Detection of S-phase cells by EdU staining. (ii) *RAD51* mRNA detection by smFISH. Scale bars, 20 μm. (**c**) *RAD51* mRNA and EdU staining images of 10 μM zeocin sample with higher magnification showing *RAD51* transcripts on EdU-negative cells. (i) Merged image showing *RAD51* mRNA and EdU signals. White dashed box delineates magnified area. Scale bar, 20 μm. (ii) Magnified area showing *RAD51* mRNA signals. Scale bar, 5 μm. (iii) Magnified area showing EdU signals. Scale bar, 5 μm. (**d**) Percentage of EdU positive cells in roots after exposure to 0 μM, 10 μM, 50 μM, 170 μM zeocin. ANOVA revealed statistically significant difference in EdU positive cell numbers by zeocin concentration (F(3)=24.38, *p*=1.67e-08)) in our measurements (n=1123, n=1356, n=1519, n=1208 measurements for 0 μM, 10 μM, 50 μM, 170 μM zeocin respectively). Error bars indicate standard deviation. Letters indicate results of TukeyHSD test with 95% confidence level. (**e**) Schematic representation of the cell cycle and its phases. Cell cycle phases labelled by corresponding fluorescent reporter indicated for each of the plant lines used for the cell cycle analysis. (**f**) Representation of cells in different phases of the cell cycle in Cytrap line roots, value shown in %. Representation of G1 cells is an approximation calculated by exclusion according to Cytrap line description. Dataset containing 1002, 816, 800, 998 individual measurements for 0 μM, 50 μM and 170 μM concentrations correspondingly was used. (**g**) Percentage of G1-phase cells in roots after for the different concentrations of zeocin in roots of PlaCCI plant line. ANOVA revealed statistically significant difference in EdU G1-phase cell numbers by zeocin concentration (F(3)=15.15, *p*=4.56e-07)) in our measurements (n=2526, n=2438, n=1997, n=1698 measurements for 0 μM, 10 μM, 50 μM, 170 μM zeocin respectively). Error bars indicate standard deviation. Letters indicate results of TukeyHSD test with 95% confidence level. (**h**) Representative confocal image of root cells from CDT1a-CFP plant line after exposure to 50 μM zeocin, showing with *RAD51* mRNA signal detection via WM-smFISH. (i) Detection of CDT1a-CFP reporter. (ii) Detection *RAD51* mRNA signals. Asterisks indicate transcription sites. (iii) Overlay of (i) and (ii) images. Scale bars, 20 μm.

To further investigate the association between *RAD51* transcription and S/G2 phases of the cell cycle under damage, we used Cytrap (Yin *et al*., 2014), CDT1-CFP (Desvoyes *et al*., 2019) and PlaCCI (Desvoyes *et al*., 2020) lines, which express fluorescent reporters specific to individual cell cycle phases. Cytrap line allows visualization of S/G2 phase cells and G2/M cells while PlaCCI line provides additional possibility of direct G1 phase cells visualization using CDT1a-CFP construct which is also available as a separate line (**Fig. 4e**). Analysis of Cytrap line revealed a decrease in S/G2-phase cells with increasing concentrations of zeocin, consistent with EdU staining data (**Fig. S8a**), as well as an increase in the fraction of cells expressing G2/M reporter, potentially corresponding to checkpoint arrest (**Fig. S8b**) (Preuss & Britt, 2003). Statistical analysis revealed that the changes in S/G2 and G2/M phase cells can be explained by both zeocin concentration and the cell type with significant interaction between the two parameters (*p* = 1.08e^-09^, *p* = 6.56e^-06^ accordingly) (**Fig. S8e-f**). Further pairwise comparisons revealed statistically significant changes between the concentrations only within root stele group, an observation strikingly similar to the earlier reported *RAD51* transcriptional data. Of note, the combined percentage of S/G2 and G2/M cells at higher concentrations of zeocin suggests that a large fraction of cells is not either in S phase or at G2/M checkpoint, thus potentially residing in G1 phase (**Fig. 4f**). Considering the novelty of this finding we decided to rely on direct visualization of G1 phase cells using recently developed PlaCCI and CDT1-CFP lines. The results confirmed an increase in the amount of G1 cells in response to increasing concentrations of zeocin in PlaCCI and CDT1-CFP lines indicating potential cell cycle arrest at G1/S checkpoint (**Fig. 4g, Fig. S9a**). The proportion of cells with G2/M marker also increased confirming the results obtained with Cytrap line (**Fig. S9b, S9c**). Analysis of the results showed the changes in G1 cell numbers can be explained by both zeocin concentration and the cell type with significant interaction between the two parameters (*p* = 0.0268) (**Fig. S9a, S9d**). Further pairwise comparisons revealed statistically significant increase in G1 cells only within root stele group.

Intriguingly, we also observed a small fraction of cells without any fluorescent reporter presence in roots of PlaCCI line (**Fig. S9c**).

One possible explanation for *RAD51* transcripts observed in G1 cells is that they could be produced in S/G2 and carried over to G1 due to a potentially long half-life of transcripts. To evaluate the mRNA half-life, we treated seedlings with the transcription elongation inhibitor, actinomycin D (ActD), and conducted a time-series smFISH analysis (**Fig. S10**). The half-life of the *RAD51* mRNA was calculated from our data as 4.9 hours. Considering this measurement there is the possibility of *RAD51* mRNA persisting beyond the G2-phase of the cell cycle. Indeed, we detected mitotic cells, which normally do not actively transcribe genes, possessing *RAD51* mRNAs (**Fig. S10c**). However, this half-life (4.9h) is relatively short compared to the cell cycle duration (Rahni & Birnbaum, 2019) so while *RAD51* mRNA may be carried between cell divisions, its half-life alone seems unlikely to explain the high proportion of cells with *RAD51* mRNA signals in zeocin samples.

To show *RAD51* transcription in G1 arrested cells directly, we performed *RAD51* smFISH detection on CDT1-CFP line, expressing the same G1 marker as PlaCCI line alone (**Fig. 4h**). The results clearly show the presence of multiple *RAD51* mRNAs and most importantly active transcriptional sites, as judged by the presence of larger smFISH foci in the nucleus (**Fig. 4h(ii)**), in cells labeled with G1 phase reporter (**Fig. 4h(iii)**), thereby directly confirming predicted *RAD51* transcription during G1 phase under DNA damage. Upon further examination of transcription site numbers in G1 and non-G1 cells using the CDT1-CFP line, we observed a nearly equal partitioning (Supplementary table 1). This observation implies that *RAD51* transcription occurs with approximately equal probability during both G1 and other stages of the cell cycle under conditions of DNA damage. This result correlates with similar numbers of cells residing in G1 and other cell cycle phases (**Fig. 4f, S9c**). Intriguingly, some G1 *RAD51* transcription was possible even under control conditions.

## DISCUSSION

This study describes the transcriptional activation of *RAD51* following increasing amounts of DNA damage. Our findings indicate a rise in total mRNA production that results from an increase in transcriptional output per cell as DNA damage increases. These results underscore the cell’s capacity to sense the extent of damage and modulate *RAD51* transcription accordingly.

Using single-cell measurements by smFISH technique we obtained data showing differences in DNA damage sensitivity between cells, manifested by varying *RAD51* mRNA transcriptional output in response to the same concentration of DNA damaging agent. Our results also revealed dynamic changes in numbers of cells residing in different cell cycle phases in response to increased DNA damage. Of note, these changes were not as evident between samples treated with 50 μM and 170 μM zeocin concentrations, for which the number of *RAD51* mRNAs per cell did not differ substantially. One can therefore speculate that proportions of cell arrested at different cell cycle phases achieved in 50 μM and 170 μM zeocin treated samples are the most efficient for cells to cope with DNA damage.

Cell cycle arrest and upregulation of DDR genes are the two key elements of the DDR response. Our data from several independent experiments showed that root stele cells consistently differed from other cell types in both cell cycle changes and *RAD51* transcriptional response to growing amounts of DNA damage. Specifically, root stele cells exhibited a more extensive *RAD51* transcriptional activation as well as larger fluctuations in numbers of cells represented at different cell cycle stages under the same zeocin concentrations. One potential explanation for this observation could be linked to distinct cell cycle duration among the different cell types. Live-imaging experiments indicate a shorter cell cycle duration for stele cells (∼15 hours) compared to other root cell types (∼23 hours for cortex and 24h for the epidermis) (Rahni & Birnbaum, 2019). Faster proliferation rates have been correlated with increased susceptibility to DNA damage (Kiraly *et al*., 2015; Alhmoud *et al*., 2020). Notably, stele cells have demonstrated higher sensitivity to cell death in response to zeocin (Yoshiyama *et al*., 2017; Johnson *et al*., 2018; Ryu *et al*., 2019). Consequently, the greater accumulation of damage may underscore the elevated transcriptional response of *RAD51* in stele cells.

The observation of *RAD51* transcription occurring outside S/G2 phases of the cell cycle is another important finding of this study. DDR via HR and *RAD51* gene expression has been associated with S/G2-phase of the cell cycle in many organisms (Basile *et al*., 1992; Yamamoto *et al*., 1996; Doutriaux *et al*., 1998). In *A. thaliana RAD51* transcription in response to DNA damage was coincident with the cell cycle arrest at G2/M checkpoint (Osakabe *et al*., 2005; De Schutter *et al*., 2007). Later studies demonstrated that both G1/S and G2/M checkpoints can be used to ensure cell cycle arrest in response to DNA damage. For example, hydroxyurea (HU) treatment was shown to activate both G2/M cell cycle arrest (De Schutter *et al*., 2007) and G1/S checkpoint (Saban & Bujak, 2009; Cabral *et al*., 2020), a phenomenon also observed in response to gamma irradiation (Hefner, 2003; Hefner *et al*., 2006; Ricaud *et al*., 2007). Zeocin, the radiomimetic drug used in this study to induce DSBs, was so far reported to promote arrest at the G2/M checkpoint (De Schutter *et al*., 2007; Chen *et al*., 2017). Our findings challenge this view by demonstrating that a considerable number of cells undergo arrest in the G1 phase while still exhibiting *RAD51* transcription. This observation does not mean that *RAD51* is not transcribed in S/G2. Indeed, our data indicated equal representation of transcription site signals in G1 cells and cells in the rest of the cell cycle, potentially indicating absence of preference for *RAD51* transcription between the cell cycle phases under varying amounts of DNA damage. Our results therefore suggest *RAD51* transcription being more widespread across the cell cycle than initially anticipated.

Previous studies suggest a potential reason and implication behind the release of the S/G2 restriction of *RAD51* expression. For instance, it was shown that repetitive sequences can be repaired via HR during G1 phase, proven by the recruitment of RAD51 to centromeric break sites in mouse and human cells (Yilmaz *et al*., 2021). The HR machinery is also involved in G1 repair of ribosomal DNA, another type of repetitive sequence in human cell cultures (van Sluis & McStay, 2015). Moreover, non-recombinogenic functions in DNA reparation were suggested for RAD51 and some HR proteins (Cano-Linares *et al*., 2021; Prado, 2021). We suggest this as one of the potential reasons behind our observation of active *RAD51* transcription in G1 after DNA damage exposure. Also, in *A. thaliana*, RAD54 foci were shown to emerge with high frequency in both in G1 and G2 cells after gamma irradiation (Hirakawa & Matsunaga, 2019). The necessity of prior RAD51 foci formation for the formation of RAD54 foci points to the possibility of RAD51 foci presence in G1 phase *A. thaliana* cells (Hirakawa *et al*., 2017).

Altogether, the results of this article shed new light on the DNA damage response in plants, uncovering distinctions in the transcriptional response of *RAD51* across various cell types. Moreover, it highlights the noteworthy occurrence of transcription during the G1 phase of the cell cycle.

## ACKNOWLEDGEMENTS

We would like to acknowledge Lihua Zhao for advice on whole mount smFISH protocol performance. We are grateful to Benedicte Desvoyes and Crisanto Gutierrez for providing seeds of PlaCCI and CDT1-CFP lines and critical reading of the manuscript. This work was supported by Swedish Research Council (Vetenskapsrådet) grant number 2018-04101 to SR; Knut and Alice Wallenberg Foundation (KAW 2019-0062).

## AUTHOR CONTRIBUTIONS

KK, SvR, AM, AS and SR designed the research. KK performed the research and data analysis. AF assisted data analysis and created software pipelines for image analysis. KS assisted statistical analysis of the data. KK and SR wrote the manuscript. CW provided material for the study. CW and AS critically read the manuscript.

## DATA AVAILABILITY

The data from this study are not currently deposited in external repositories. However, they can be requested directly from the corresponding author. Upon acceptance, the data will be made available in repositories.

## REFERENCES

Abe K, Osakabe K, Nakayama S, Endo M, Tagiri A, Todoriki S, Ichikawa H, Toki S. 2005. Arabidopsis RAD51C gene is important for homologous recombination in meiosis and mitosis. Plant Physiology 139: 896–908.

Adachi S, Minamisawa K, Okushima Y, Inagaki S, Yoshiyama K, Kondou Y, Kaminuma E, Kawashima M, Toyoda T, Matsui M, et al. 2011. Programmed induction of endoreduplication by DNA double-strand breaks in *Arabidopsis*. Proceedings of the National Academy of Sciences 108: 10004–10009.

Aki SS, Umeda M. 2016. Cytrap Marker Systems for In Vivo Visualization of Cell Cycle Progression in Arabidopsis. In: Caillaud M-C, ed. Methods in Molecular Biology. Plant Cell Division. New York, NY: Springer New York, 51–57.

Alhmoud JF, Woolley JF, Al Moustafa A-E, Malki MI. 2020. DNA Damage/Repair Management in Cancers. Cancers 12: 1050.

Banerjee S, Roy S. 2021. An insight into understanding the coupling between homologous recombination mediated DNA repair and chromatin remodeling mechanisms in plant genome: an update. *Cell Cycle (Georgetown*, Tex.) 20: 1760–1784.

Basile G, Aker M, Mortimer RK. 1992. Nucleotide sequence and transcriptional regulation of the yeast recombinational repair gene RAD51. Molecular and Cellular Biology 12: 3235– 3246.

Bee L, Fabris S, Cherubini R, Mognato M, Celotti L. 2013. The Efficiency of Homologous Recombination and Non-Homologous End Joining Systems in Repairing Double-Strand Breaks during Cell Cycle Progression (S Cotterill, Ed.). PLoS ONE 8: e69061.

Biedermann S, Harashima H, Chen P, Heese M, Bouyer D, Sofroni K, Schnittger A. 2017. The retinoblastoma homolog RBR 1 mediates localization of the repair protein RAD 51 to DNA lesions in *Arabidopsis*. The EMBO Journal 36: 1279–1297.

Bonilla B, Hengel SR, Grundy MK, Bernstein KA. 2020. *RAD51* Gene Family Structure and Function. Annual Review of Genetics 54: 25–46.

Cabral D, Banora MY, Antonino JD, Rodiuc N, Vieira P, Coelho RR, Chevalier C, Eekhout T, Engler G, De Veylder L, et al. 2020. The plant WEE1 kinase is involved in checkpoint control activation in nematode-induced galls. New Phytologist 225: 430–447.

Cano-Linares MI, Yáñez-Vilches A, García-Rodríguez N, Barrientos-Moreno M, González-Prieto R, San-Segundo P, Ulrich HD, Prado F. 2021. Non-recombinogenic roles for Rad52 in translesion synthesis during DNA damage tolerance. EMBO reports 22: e50410.

Chabot T, Defontaine A, Marquis D, Renodon-Corniere A, Courtois E, Fleury F, Cheraud Y. 2019. New Phosphorylation Sites of Rad51 by c-Met Modulates Presynaptic Filament Stability. Cancers 11: 413.

Charbonnel C, Gallego ME, White CI. 2010. Xrcc1-dependent and Ku-dependent DNA double-strand break repair kinetics in Arabidopsis plants: Double-strand break repair kinetics in Arabidopsis. The Plant Journal 64: 280–290.

Chen P, Takatsuka H, Takahashi N, Kurata R, Fukao Y, Kobayashi K, Ito M, Umeda M. 2017. Arabidopsis R1R2R3-Myb proteins are essential for inhibiting cell division in response to DNA damage. Nature Communications 8: 635.

Coïc E, Martin J, Ryu T, Tay SY, Kondev J, Haber JE. 2011. Dynamics of Homology Searching During Gene Conversion in *Saccharomyces cerevisiae* Revealed by Donor Competition. Genetics 189: 1225–1233.

Cools T, Iantcheva A, Weimer AK, Boens S, Takahashi N, Maes S, Van den Daele H, Van Isterdael G, Schnittger A, De Veylder L. 2011. The *Arabidopsis thaliana* Checkpoint Kinase WEE1 Protects against Premature Vascular Differentiation during Replication Stress. The Plant Cell 23: 1435–1448.

Cui W, Wang H, Song J, Cao X, Rogers HJ, Francis D, Jia C, Sun L, Hou M, Yang Y, et al. 2017. Cell cycle arrest mediated by Cd-induced DNA damage in Arabidopsis root tips. Ecotoxicology and Environmental Safety 145: 569–574.

Culligan K, Tissier A, Britt A. 2004. ATR Regulates a G2-Phase Cell-Cycle Checkpoint in *Arabidopsis thaliana*. The Plant Cell 16: 1091–1104.

Da Ines O, Bazile J, Gallego ME, White CI. 2022. DMC1 attenuates RAD51-mediated recombination in Arabidopsis (IR Henderson, Ed.). PLOS Genetics 18: e1010322.

Da Ines O, Degroote F, Goubely C, Amiard S, Gallego ME, White CI. 2013. Meiotic Recombination in Arabidopsis Is Catalysed by DMC1, with RAD51 Playing a Supporting Role (FCH Franklin, Ed.). PLoS Genetics 9: e1003787.

De Schutter K, Joubès J, Cools T, Verkest A, Corellou F, Babiychuk E, Van Der Schueren E, Beeckman T, Kushnir S, Inzé D, et al. 2007. Arabidopsis WEE1 kinase controls cell cycle arrest in response to activation of the DNA integrity checkpoint. The Plant Cell 19: 211–225.

Desvoyes B, Arana-Echarri A, Barea MD, Gutierrez C. 2020. A comprehensive fluorescent sensor for spatiotemporal cell cycle analysis in Arabidopsis. Nature Plants 6: 1330–1334.

Desvoyes B, Noir S, Masoud K, López MI, Genschik P, Gutierrez C. 2019. FBL17 targets CDT1a for degradation in early S-phase to prevent Arabidopsis genome instability. Plant Biology.

Doutriaux M-P, Couteau F, Bergounioux C, White C. 1998. Isolation and characterisation of the RAD51 and DMC1 homologs from Arabidopsis thaliana. Molecular and General Genetics MGG 257: 283–291.

Duncan S, Olsson TSG, Hartley M, Dean C, Rosa S. 2016. A method for detecting single mRNA molecules in Arabidopsis thaliana. Plant Methods 12: 13.

Duncan S, Olsson T, Hartley M, Dean C, Rosa S. 2017. Single Molecule RNA FISH in Arabidopsis Root Cells. BIO-PROTOCOL 7.

Fan T, Kang H, Wu D, Zhu X, Huang L, Wu J, Zhu Y. 2022. Arabidopsis γ-H2A.X-INTERACTING PROTEIN participates in DNA damage response and safeguards chromatin stability. Nature Communications 13: 7942.

Feng W, Hale CJ, Over RS, Cokus SJ, Jacobsen SE, Michaels SD. 2017. Large-scale heterochromatin remodeling linked to overreplication-associated DNA damage. Proceedings of the National Academy of Sciences 114: 406–411.

Flott S, Kwon Y, Pigli YZ, Rice PA, Sung P, Jackson SP. 2011. Regulation of Rad51 function by phosphorylation. EMBO reports 12: 833–839.

Friesner JD, Liu B, Culligan K, Britt AB. 2005. Ionizing Radiation–dependent γ-H2AX Focus Formation Requires Ataxia Telangiectasia Mutated and Ataxia Telangiectasia Mutated and Rad3-related. Molecular Biology of the Cell 16: 2566–2576.

Gamborg OL, Murashige T, Thorpe TA, Vasil IK. 1976. Plant tissue culture media. In Vitro 12: 473–478.

Goldfarb T, Lichten M. 2010. Frequent and Efficient Use of the Sister Chromatid for DNA Double-Strand Break Repair during Budding Yeast Meiosis (RS Hawley, Ed.). PLoS Biology 8: e1000520.

Gong Z-Y, Kidoya H, Mohri T, Han Y, Takakura N. 2014. DNA Damage Enhanced by the Attenuation of SLD5 Delays Cell Cycle Restoration in Normal Cells but Not in Cancer Cells (R Morishita, Ed.). PLoS ONE 9: e110483.

Hefner E. 2003. Arabidopsis mutants sensitive to gamma radiation include the homologue of the human repair gene ERCC1. Journal of Experimental Botany 54: 669–680.

Hefner E, Huefner N, Britt AB. 2006. Tissue-specific regulation of cell-cycle responses to DNA damage in Arabidopsis seedlings. DNA Repair 5: 102–110.

Hicks WM, Yamaguchi M, Haber JE. 2011. Real-time analysis of double-strand DNA break repair by homologous recombination. Proceedings of the National Academy of Sciences 108: 3108–3115.

Hirakawa T, Hasegawa J, White CI, Matsunaga S. 2017. RAD 54 forms DNA repair foci in response to DNA damage in living plant cells. The Plant Journal 90: 372–382.

Hirakawa T, Matsunaga S. 2019. Characterization of DNA Repair Foci in Root Cells of Arabidopsis in Response to DNA Damage. Frontiers in Plant Science 10: 990.

Johnson RD. 2000. Sister chromatid gene conversion is a prominent double-strand break repair pathway in mammalian cells. The EMBO Journal 19: 3398–3407.

Johnson RA, Conklin PA, Tjahjadi M, Missirian V, Toal T, Brady SM, Britt AB. 2018. SUPPRESSOR OF GAMMA RESPONSE1 Links DNA Damage Response to Organ Regeneration. Plant Physiology 176: 1665–1675.

Kiraly O, Gong G, Olipitz W, Muthupalani S, Engelward BP. 2015. Inflammation-Induced Cell Proliferation Potentiates DNA Damage-Induced Mutations In Vivo (R Risques, Ed.). PLOS Genetics 11: e1004901.

Lang J, Smetana O, Sanchez-Calderon L, Lincker F, Genestier J, Schmit A, Houlné G, Chabouté M. 2012. Plant γH2AX foci are required for proper DNA DSB repair responses and colocalize with E2F factors. New Phytologist 194: 353–363.

Lee Y, Wang Q, Shuryak I, Brenner DJ, Turner HC. 2019. Development of a high-throughput γ-H2AX assay based on imaging flow cytometry. Radiation Oncology 14: 150.

Li W, Chen C, Markmann-Mulisch U, Timofejeva L, Schmelzer E, Ma H, Reiss B. 2004. The Arabidopsis AtRAD51 gene is dispensable for vegetative development but required for meiosis. Proceedings of the National Academy of Sciences of the United States of America 101: 10596–10601.

Lim G, Chang Y, Huh W-K. 2020. Phosphoregulation of Rad51/Rad52 by CDK1 functions as a molecular switch for cell cycle–specific activation of homologous recombination. Science Advances 6: eaay2669.

Lim D-S, Hasty P. 1996. A Mutation in Mouse *rad51* Results in an Early Embryonic Lethal That Is Suppressed by a Mutation in *p53*. Molecular and Cellular Biology 16: 7133–7143.

Löbrich M, Shibata A, Beucher A, Fisher A, Ensminger M, Goodarzi AA, Barton O, Jeggo PA. 2010. γH2AX foci analysis for monitoring DNA double-strand break repair: Strengths, limitations and optimization. Cell Cycle 9: 662–669.

Masson J-Y, Stasiak AZ, Stasiak A, Benson FE, West SC. 2001. Complex formation by the human RAD51C and XRCC3 recombination repair proteins. Proceedings of the National Academy of Sciences 98: 8440–8446.

Meschichi A, Zhao L, Reeck S, White C, Da Ines O, Sicard A, Pontvianne F, Rosa S. 2022. The plant-specific DDR factor SOG1 increases chromatin mobility in response to DNA damage. EMBO reports 23: e54736.

Mueller F, Senecal A, Tantale K, Marie-Nelly H, Ly N, Collin O, Basyuk E, Bertrand E, Darzacq X, Zimmer C. 2013. FISH-quant: automatic counting of transcripts in 3D FISH images. Nature Methods 10: 277–278.

Muschel RJ, Zhang HB, Iliakis G, McKenna WG. 1991. Cyclin B expression in HeLa cells during the G2 block induced by ionizing radiation. Cancer Research 51: 5113–5117.

Musielak TJ, Schenkel L, Kolb M, Henschen A, Bayer M. 2015. A simple and versatile cell wall staining protocol to study plant reproduction. Plant Reproduction 28: 161–169.

Narsai R, Howell KA, Millar AH, O’Toole N, Small I, Whelan J. 2007. Genome-Wide Analysis of mRNA Decay Rates and Their Determinants in *Arabidopsis thaliana*. The Plant Cell 19: 3418–3436.

Osakabe K, Abe K, Yamanouchi H, Takyuu T, Yoshioka T, Ito Y, Kato T, Tabata S, Kurei S, Yoshioka Y, et al. 2005. Arabidopsis Rad51B is important for double-strand DNA breaks repair in somatic cells. Plant Molecular Biology 57: 819–833.

Prado F. 2021. Non-Recombinogenic Functions of Rad51, BRCA2, and Rad52 in DNA Damage Tolerance. Genes 12: 1550.

Preuss SB, Britt AB. 2003. A DNA-Damage-Induced Cell Cycle Checkpoint in Arabidopsis. Genetics 164: 323–334.

Rahni R, Birnbaum KD. 2019. Week-long imaging of cell divisions in the Arabidopsis root meristem. Plant Methods 15: 30.

Raleigh JM, O’Connell MJ. 2000. The G2 DNA damage checkpoint targets both Wee1 and Cdc25. Journal of Cell Science 113: 1727–1736.

Redon CE, Dickey JS, Bonner WM, Sedelnikova OA. 2009. γ-H2AX as a biomarker of DNA damage induced by ionizing radiation in human peripheral blood lymphocytes and artificial skin. Advances in Space Research 43: 1171–1178.

Ricaud L, Proux C, Renou J-P, Pichon O, Fochesato S, Ortet P, Montané M-H. 2007. ATM-Mediated Transcriptional and Developmental Responses to γ-rays in Arabidopsis (S Kepinski, Ed.). PLoS ONE 2: e430.

Rogakou EP, Boon C, Redon C, Bonner WM. 1999. Megabase Chromatin Domains Involved in DNA Double-Strand Breaks in Vivo. The Journal of Cell Biology 146: 905–916.

Ryu TH, Go YS, Choi SH, Kim J, Chung BY, Kim J. 2019. SOG 1-dependent NAC 103 modulates the DNA damage response as a transcriptional regulator in Arabidopsis. The Plant Journal 98: 83–96.

Saban N, Bujak M. 2009. Hydroxyurea and hydroxamic acid derivatives as antitumor drugs. Cancer Chemotherapy and Pharmacology 64: 213–221.

Saintigny Y, Delacôte F, Boucher D, Averbeck D, Lopez BS. 2007. XRCC4 in G1 suppresses homologous recombination in S/G2, in G1 checkpoint-defective cells. Oncogene 26: 2769– 2780.

Saleh-Gohari N. 2004. Conservative homologous recombination preferentially repairs DNA double-strand breaks in the S phase of the cell cycle in human cells. Nucleic Acids Research 32: 3683–3688.

Schuermann D, Molinier J, Fritsch O, Hohn B. 2005. The dual nature of homologous recombination in plants. Trends in Genetics 21: 172–181.

Shinohara A, Ogawa H, Ogawa T. 1992. Rad51 protein involved in repair and recombination in S. cerevisiae is a RecA-like protein. Cell 69: 457–470.

van Sluis M, McStay B. 2015. A localized nucleolar DNA damage response facilitates recruitment of the homology-directed repair machinery independent of cell cycle stage. Genes & Development 29: 1151–1163.

Sørensen CS, Hansen LT, Dziegielewski J, Syljuåsen RG, Lundin C, Bartek J, Helleday T. 2005. The cell-cycle checkpoint kinase Chk1 is required for mammalian homologous recombination repair. Nature Cell Biology 7: 195–201.

Sorenson RS, Deshotel MJ, Johnson K, Adler FR, Sieburth LE. 2018. *Arabidopsis* mRNA decay landscape arises from specialized RNA decay substrates, decapping-mediated feedback, and redundancy. Proceedings of the National Academy of Sciences 115.

Stewart GS, Wang B, Bignell CR, Taylor AMR, Elledge SJ. 2003. MDC1 is a mediator of the mammalian DNA damage checkpoint. Nature 421: 961–966.

Stirling DR, Swain-Bowden MJ, Lucas AM, Carpenter AE, Cimini BA, Goodman A. 2021. CellProfiler 4: improvements in speed, utility and usability. BMC Bioinformatics 22: 433.

Su H, Cheng Z, Huang J, Lin J, Copenhaver GP, Ma H, Wang Y. 2017. Arabidopsis RAD51, RAD51C and XRCC3 proteins form a complex and facilitate RAD51 localization on chromosomes for meiotic recombination. PLoS genetics 13: e1006827.

Tsuzuki T, Fujii Y, Sakumi K, Tominaga Y, Nakao K, Sekiguchi M, Matsushiro A, Yoshimura Y, Morita T. 1996. Targeted disruption of the Rad51 gene leads to lethality in embryonic mice. Proceedings of the National Academy of Sciences 93: 6236–6240.

Vítor AC, Huertas P, Legube G, de Almeida SF. 2020. Studying DNA Double-Strand Break Repair: An Ever-Growing Toolbox. Frontiers in Molecular Biosciences 7: 24.

Wang S, Durrant WE, Song J, Spivey NW, Dong X. 2010. *Arabidopsis* BRCA2 and RAD51 proteins are specifically involved in defense gene transcription during plant immune responses. Proceedings of the National Academy of Sciences 107: 22716–22721.

Wang Y, Xiao R, Wang H, Cheng Z, Li W, Zhu G, Wang Y, Ma H. 2014. The Arabidopsis RAD51 paralogs RAD51B, RAD51D and XRCC2 play partially redundant roles in somatic DNA repair and gene regulation. The New Phytologist 201: 292–304.

Weimer AK, Biedermann S, Harashima H, Roodbarkelari F, Takahashi N, Foreman J, Guan Y, Pochon G, Heese M, Van Damme D, et al. 2016. The plant-specific CDKB 1-CYCB 1 complex mediates homologous recombination repair in *Arabidopsis*. The EMBO Journal 35: 2068–2086.

West CE, Waterworth WM, Sunderland PA, Bray CM. 2004. *Arabidopsis* DNA double-strand break repair pathways. Biochemical Society Transactions 32: 964–966.

Woo T-T, Chuang C-N, Higashide M, Shinohara A, Wang T-F. 2020. Dual roles of yeast Rad51 N-terminal domain in repairing DNA double-strand breaks. Nucleic Acids Research 48: 8474–8489.

Woo T-T, Chuang C-N, Wang T-F. 2021. Budding yeast Rad51: a paradigm for how phosphorylation and intrinsic structural disorder regulate homologous recombination and protein homeostasis. Current Genetics 67: 389–396.

Yamamoto A, Yagi H, Habu T, Yoshimura Y, Matsushiro A, Nishimune Y, Morita T, Taki T, Yoshida K, Yamamoto K, et al. 1996. Cell cycle-dependent expression of the mouseRad51 gene in proliferating cells. Molecular and General Genetics MGG 251: 1–12.

Yata K, Lloyd J, Maslen S, Bleuyard J-Y, Skehel M, Smerdon SJ, Esashi F. 2012. Plk1 and CK2 Act in Concert to Regulate Rad51 during DNA Double Strand Break Repair. Molecular Cell 45: 371–383.

Yilmaz D, Furst A, Meaburn K, Lezaja A, Wen Y, Altmeyer M, Reina-San-Martin B, Soutoglou E. 2021. Activation of homologous recombination in G1 preserves centromeric integrity. Nature 600: 748–753.

Yin K, Ueda M, Takagi H, Kajihara T, Sugamata Aki S, Nobusawa T, Umeda-Hara C, Umeda M. 2014. A dual-color marker system for *in vivo* visualization of cell cycle progression in Arabidopsis. The Plant Journal 80: 541–552.

Yoshiyama KO, Kaminoyama K, Sakamoto T, Kimura S. 2017. Increased Phosphorylation of Ser-Gln Sites on SUPPRESSOR OF GAMMA RESPONSE1 Strengthens the DNA Damage Response in *Arabidopsis thaliana*. The Plant Cell 29: 3255–3268.

Yu C, Hou L, Huang Y, Cui X, Xu S, Wang L, Yan S. 2023. The MULTI-BRCT domain protein DDRM2 promotes the recruitment of RAD51 to DNA damage sites to facilitate homologous recombination. New Phytologist 238: 1073–1084.

Zhao L, Fonseca A, Meschichi A, Sicard A, Rosa S. 2023. Whole-mount smFISH allows combining RNA and protein quantification at cellular and subcellular resolution. Nature Plants 9: 1094–1102.

