## Supplementary Figures and Tables for "Differences in *RAD51* transcriptional response and cell cycle dynamics reveal varying sensitivity to DNA damage among *Arabidopsis thaliana* root cell types"

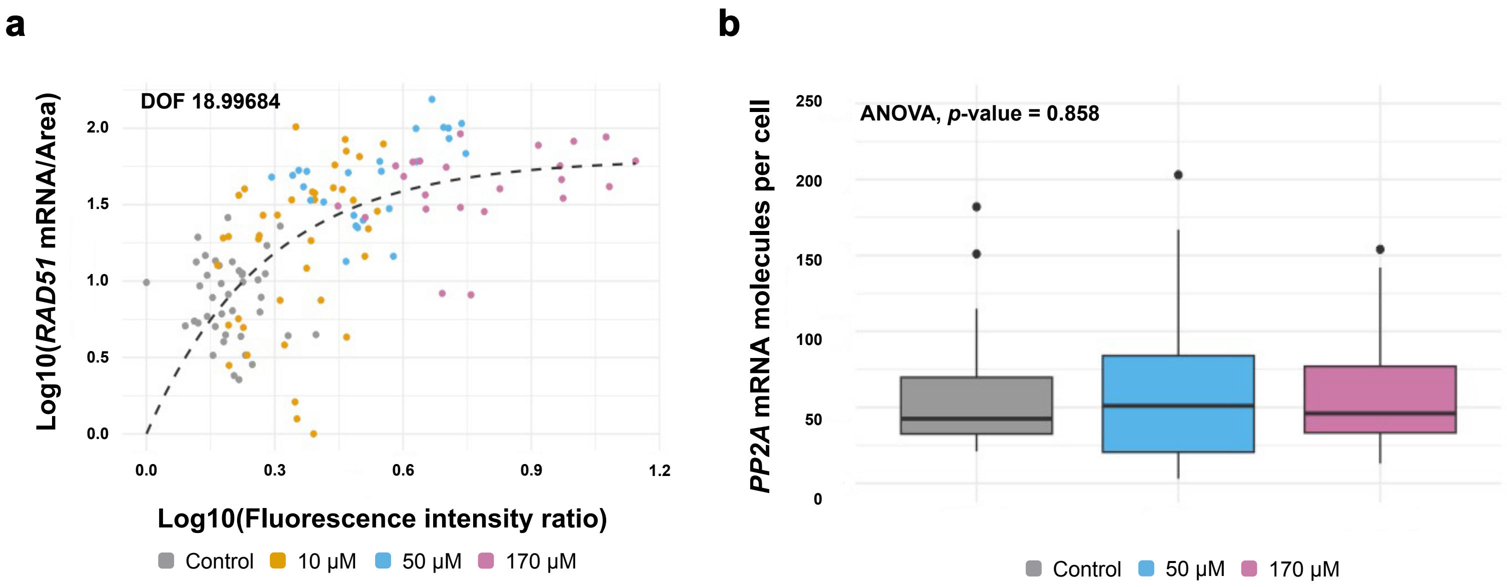
a
DOF 18.99684
Log10(RAD51 mRNA/Area)
2.0
1.5
1.0
0.5
0.0
0.0
0.3
0.6
0.9
1.2
Log10(Fluorescence intensity ratio)
Control
10 μM
50 μM
170 μM
b
ANOVA, p-value = 0.858
PP2A mRNA molecules per cell
250
200
150
100
50
0
Control
50 μM
170 μM

**Fig. S1 Evaluation of *PP2A* transcription in response to growing amounts of DNA damage in squashed roots of *Arabidopsis thaliana*.** (**a**) Correlation analysis between the number of *RAD51* mRNA molecules and ɣ-H2AX signal intensity in individual cells of squashed roots with exponential model fit. Number of *RAD51* transcripts normalized by corresponding cell area, log10 of this value used for the corresponding axis. ɣ-H2AX fluorescence intensity measured as lg(Integrated Density) with prior normalization to DAPI Integrated density. DOF indicates deviance of fit calculated for the model. Dataset contains n=40, n=40, n=25, n=22 measurements for 0 μM, 10 μM, 50 μM, 170 μM zeocin respectively. (**b**) Numbers of *PP2A* mRNA molecules per cell in samples exposed to 0 μM, 50 μM, 170 μM concentrations of zeocin. ANOVA did not reveal statistically significant difference in transcripts per cell ($F(2)=0.153$, $p=0.858$)) in our measurements (n=30, n=90, n=69 cells for 0 μM, 50 μM and 170 μM zeocin respectively).

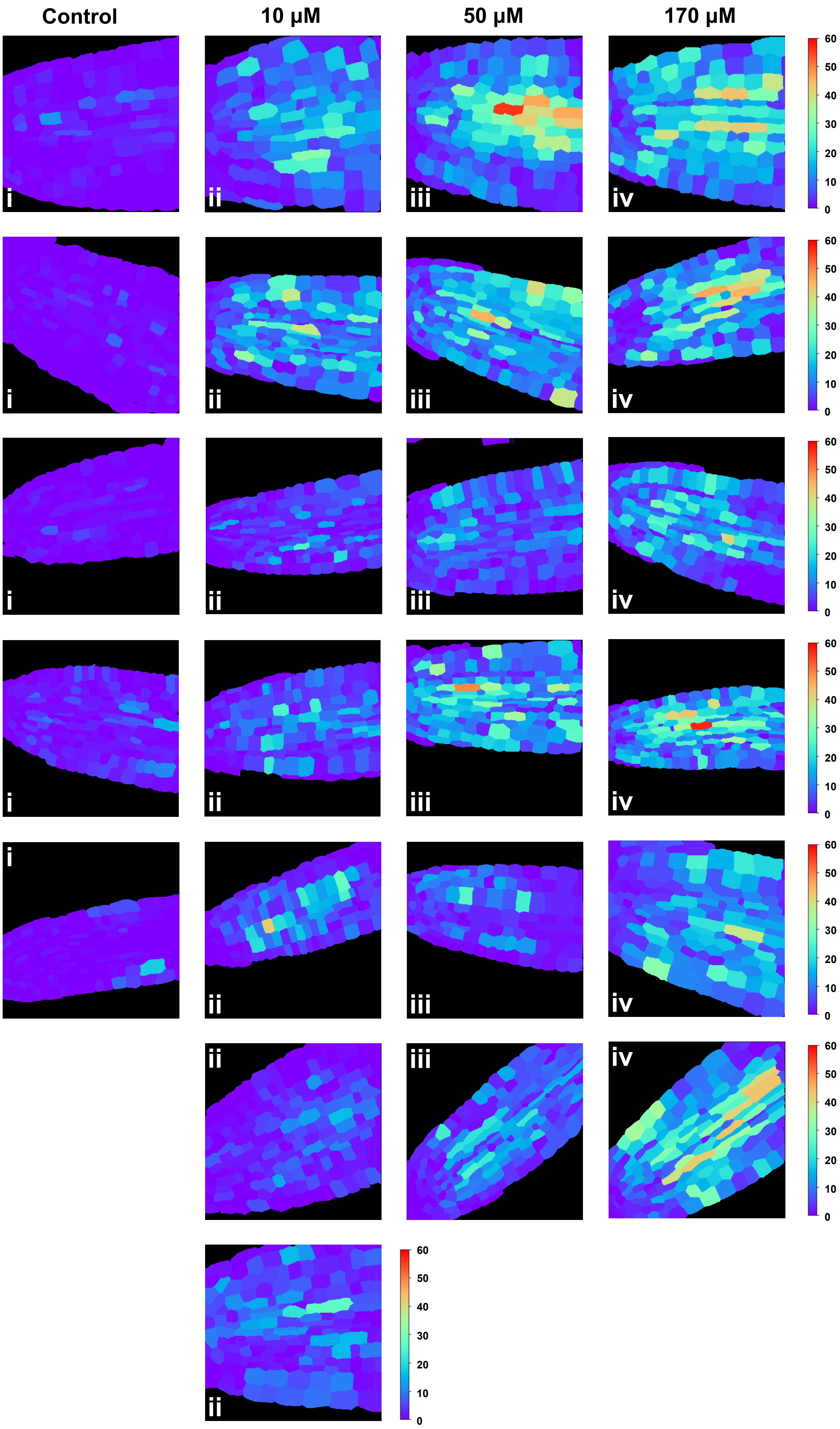
Control
10 μM
50 μM
170 μM
i
ii
iii
iv
60
50
40
30
20
10
0

**Fig. S2 Quantification of *RAD51* mRNA molecules in Col-0 *Arabidopsis thaliana* roots exposed to growing amounts of DNA damage using whole-mount smFISH.** (i, ii, iii, iv) Heatmaps represent the number of *RAD51* mRNA molecules detected in individual cells. Roots exposed to 0 μM, 10 μM, 50 μM, 170 μM concentrations of zeocin.

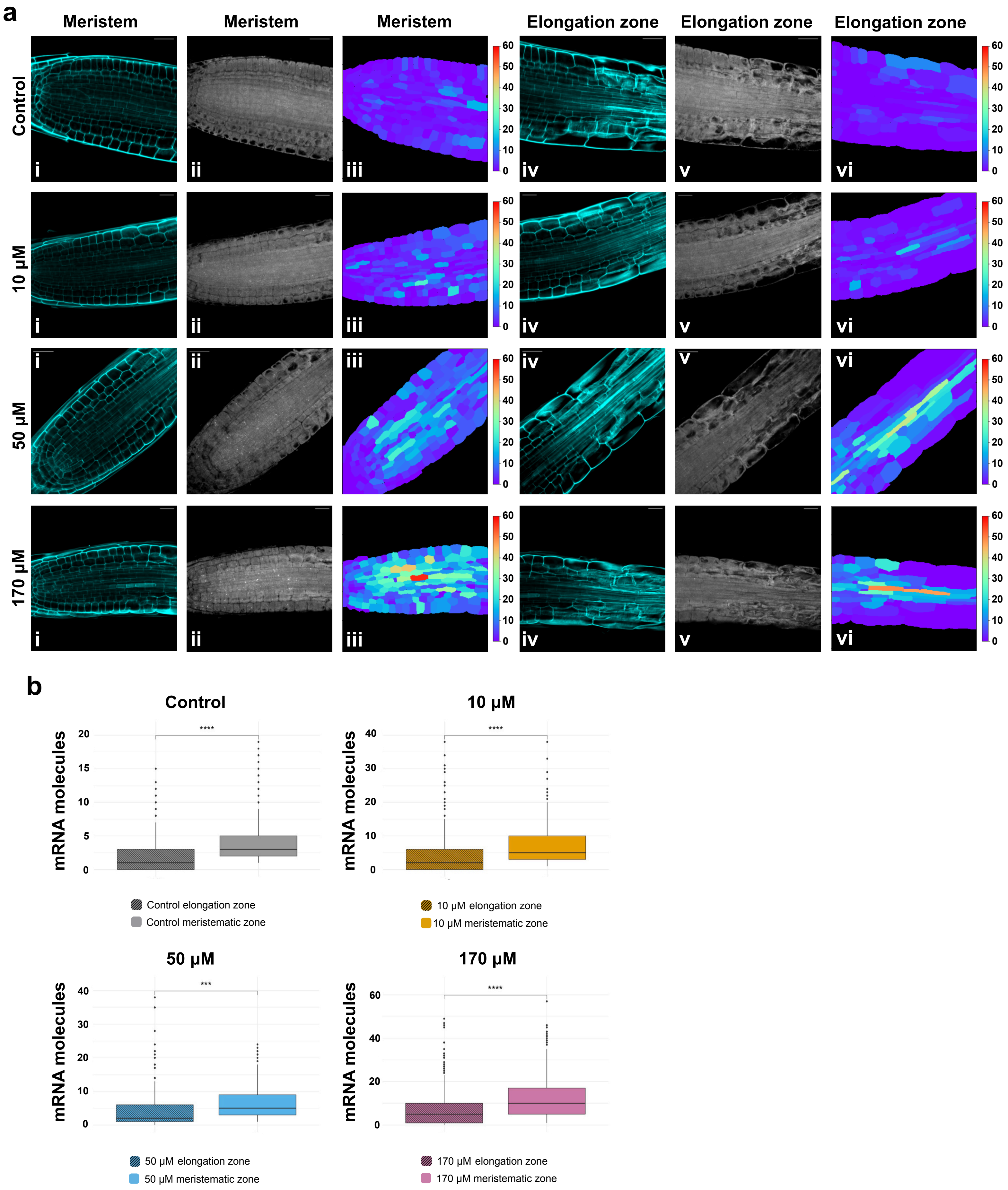
a
Meristem
Meristem
Meristem
Elongation zone
Elongation zone
Elongation zone
Control
10 μM
50 μM
170 μM
i
ii
iii
iv
v
vi
60
50
40
30
20
10
0
b
Control
10 μM
50 μM
170 μM
mRNA molecules
****
***
Control elongation zone
Control meristematic zone
10 μM elongation zone
10 μM meristematic zone
50 μM elongation zone
50 μM meristematic zone
170 μM elongation zone
170 μM meristematic zone

**Fig. S3 Evaluation of *RAD51* transcriptional response in meristematic and elongation zones of the *Arabidopsis thaliana* root using whole mount smFISH.** (**a**) Confocal images and heatmaps showing the number of *RAD51* mRNA molecules detected in individual cells. (i) Images of cell wall staining using Renaissance 2200 dye in root meristem. (ii) Images of *RAD51* mRNA transcripts in root meristem. (iii) Heatmaps representing the number of *RAD51* mRNA molecules detected in individual cells of root meristem. (iv) Images of cell wall staining using Renaissance 2200 dye in root elongation zone. (v) Images of *RAD51* mRNA transcripts in root elongation zone. (vi) Heatmaps representing the number of *RAD51* mRNA molecules detected in individual cells in root elongation zone. (**b**) Number of *RAD51* mRNA molecules per cell after exposure to 0 µM, 10 µM, 50 µM, 170 µM concentrations of zeocin in different root zones. Group means compared using T-test, asterisks show p-values. (*** and ****) indicate p values <= 0.001 and 0.0001 accordingly (n=570, n=592, n=560, n=1344 cells for 0 µM, 50 µM and 170 µM zeocin respectively).

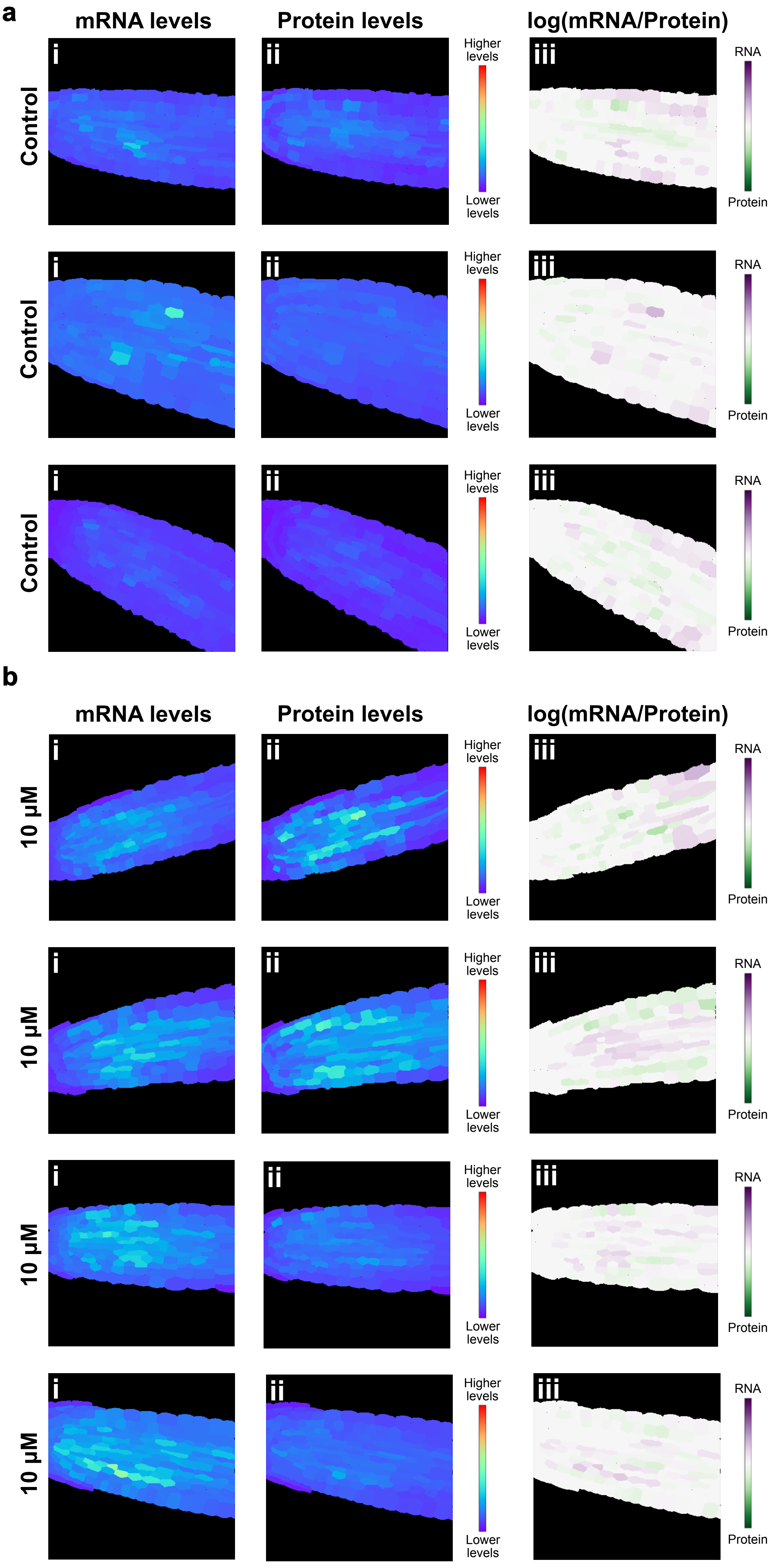
a
mRNA levels
Protein levels
log(mRNA/Protein)
Control
Control
Control
i
ii
iii
Higher levels
Lower levels
RNA
Protein
b
mRNA levels
Protein levels
log(mRNA/Protein)
10 µM
10 µM
10 µM
10 µM
i
ii
iii
Higher levels
Lower levels
RNA
Protein

**Fig. S4 Quantification of *RAD51* mRNA and RAD51-GFP protein signals in *Arabidopsis thaliana* RAD51-GFP line roots treated with 0 µM and 10 µM concentrations of zeocin.** (**a**) Roots imaged after root exposure to 0 µM and (**b**) 10 µM concentrations of zeocin. (i) Heatmaps represent the levels of the RAD51 mRNA mean signal intensity per cell. (ii) Heatmaps represent the levels of the RAD51-GFP mean signal intensity per cell. (iii) Heatmaps representing the ratio between the RNA and protein signal intensities per cell.

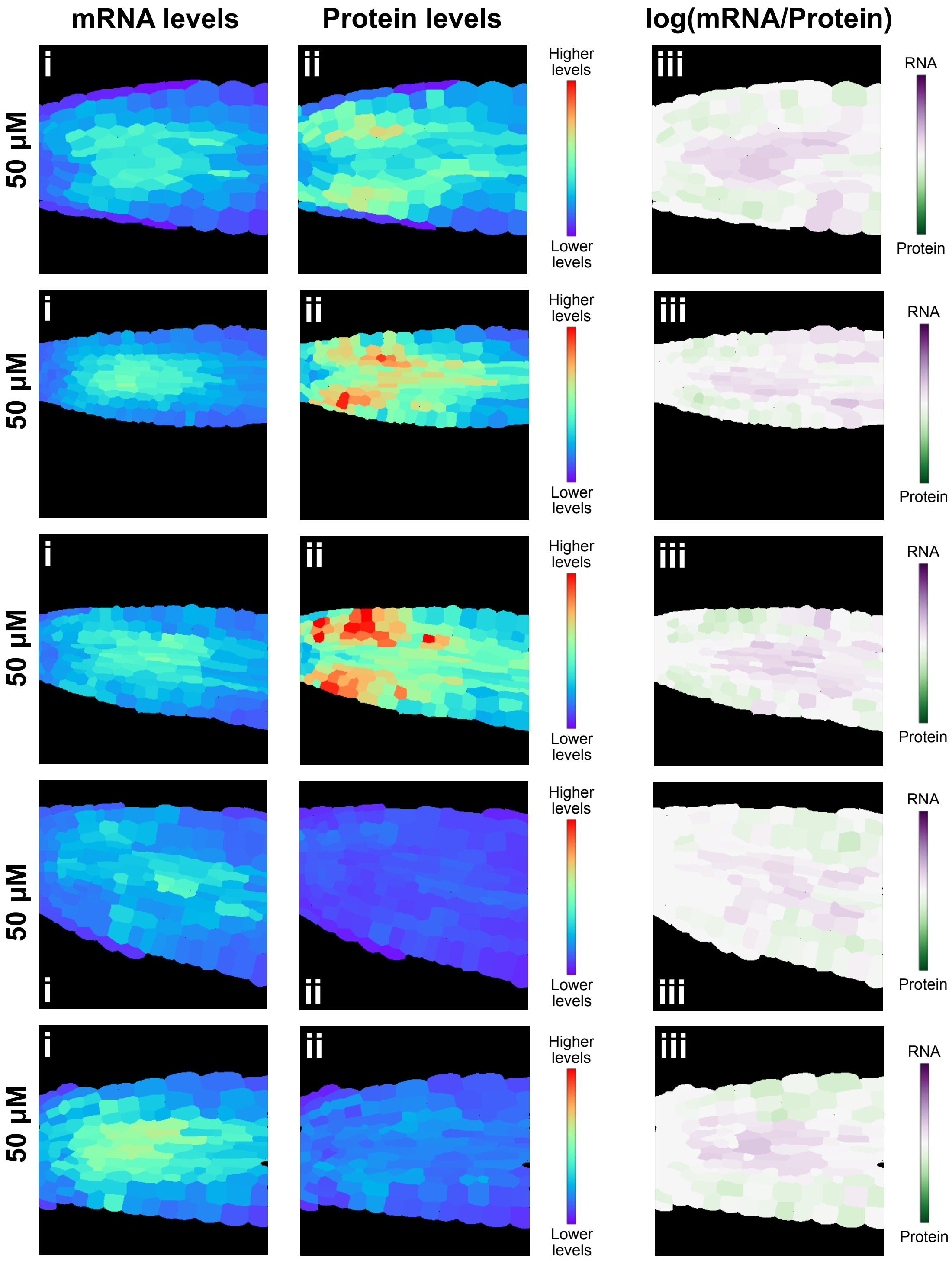
mRNA levels
Protein levels
log(mRNA/Protein)
50 µM
50 µM
50 µM
50 µM
50 µM
i
ii
iii
Higher levels
Lower levels
RNA
Protein

**Fig. S5 Quantification of *RAD51* mRNA and RAD51-GFP protein signals in *Arabidopsis thaliana* RAD51-GFP line roots treated with 50 µM concentration of zeocin.** (i, ii) Heatmaps represent the levels of the mean signal intensity per cell. (iii) Heatmaps representing the ratio between the RNA and protein signal intensities per cell.

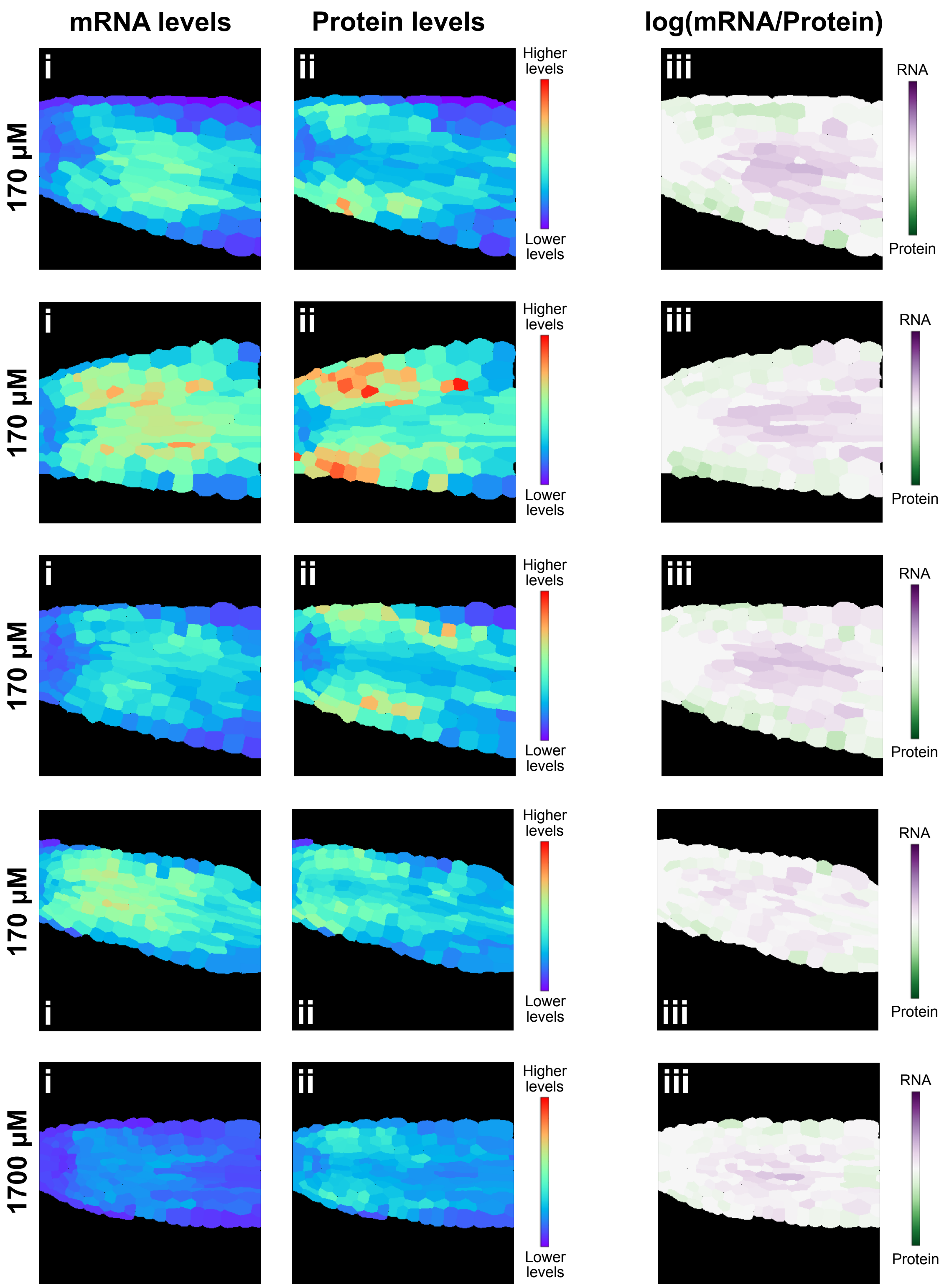

mRNA levels
Protein levels
log(mRNA/Protein)
170 µM
170 µM
170 µM
170 µM
1700 µM
i
ii
iii
Higher levels
Lower levels
RNA
Protein

**Fig. S6 Quantification of *RAD51* mRNA and RAD51-GFP protein signals in *Arabidopsis thaliana* RAD51-GFP line roots treated with 170 µM concentration of zeocin.** Signals were imaged after root exposure to 170 µM concentration of zeocin. (i, ii) Heatmaps represent the levels of the mean signal intensity per cell. (iii) Heatmaps representing the ratio between the RNA and protein signal intensities per cell.

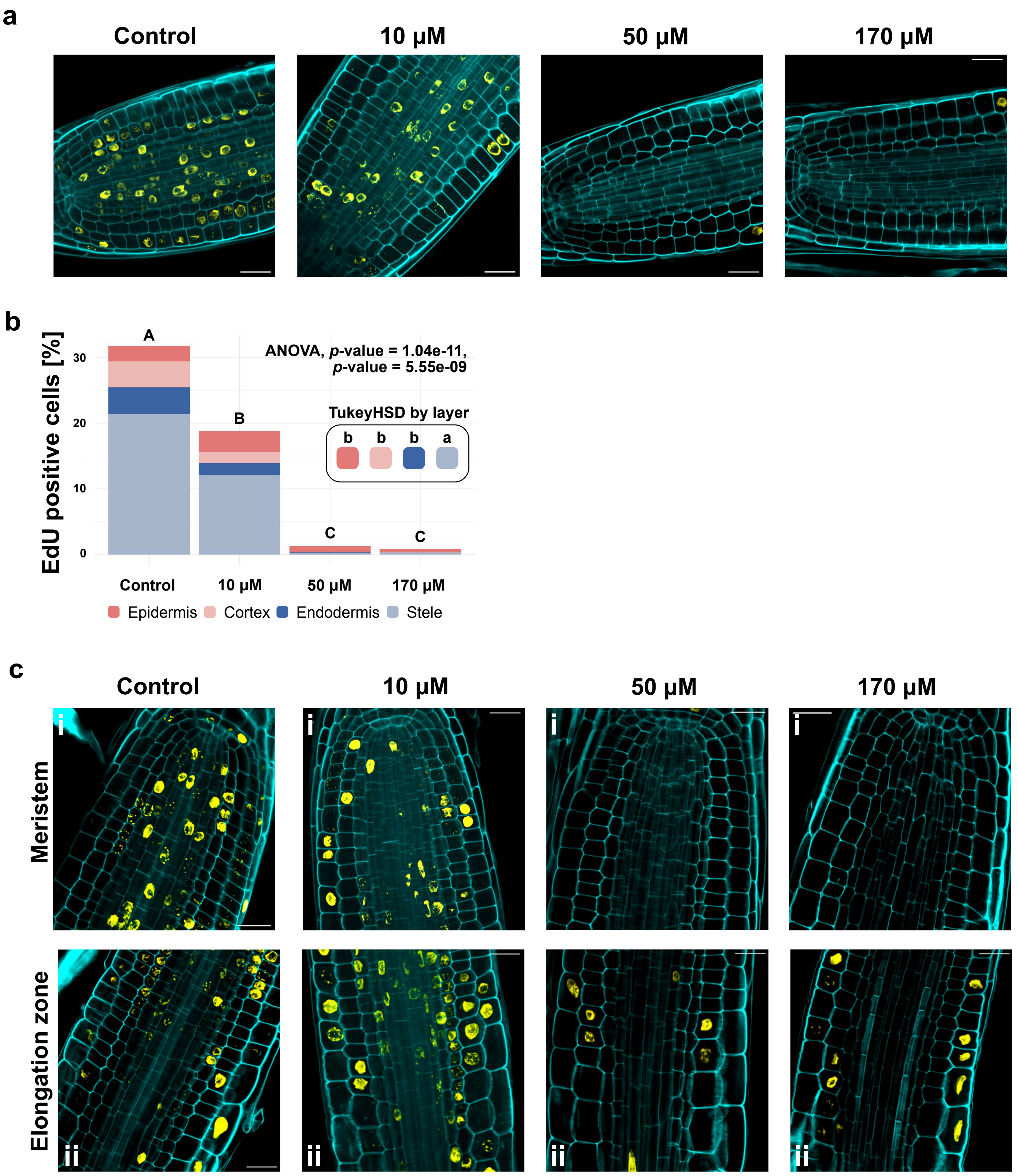
a
Control
10 µM
50 µM
170 µM
b
EdU positive cells [%]
ANOVA, p-value = 1.04e-11,
p-value = 5.55e-09
TukeyHSD by layer
b b b a
A
B
C
C
0
10
20
30
Control
10 µM
50 µM
170 µM
Epidermis
Cortex
Endodermis
Stele
c
Control
10 µM
50 µM
170 µM
Meristem
Elongation zone
i
ii

**Fig. S7 Evaluation of cell cycle arrest in *Arabidopsis thaliana* roots after exposure to growing amounts of DNA damage using EdU staining.** (**a**) Exemplary image of EdU positive cell distribution after exposure 0 μM, 10 μM, 50 μM, 170 μM concentrations of zeocin. (**b**) Evaluation of EdU positive cells relative to total number of root cells in selected cell types, value shown in %. Two-way ANOVA revealed statistically significant difference in EdU positive cell representation by both zeocin concentration ($F(3)=21.92$, $p=1.04e-11$)) and cell lineage ($F(3)=15.95$, $p=5.55e-09$)) in our measurements (n=1123, n=1356, n=1519, n=1208 cells for 0 μM, 10 μM, 50 μM, 170 μM zeocin respectively). Letters indicate results of TukeyHSD test of two-way ANOVA results with 95% confidence level. (**c**) EdU positive cells in meristematic zone (i) and elongation zone (ii) after exposure to 0 μM, 10 μM, 50 μM, 170 μM concentrations of zeocin. Scale bars, 20 μm.

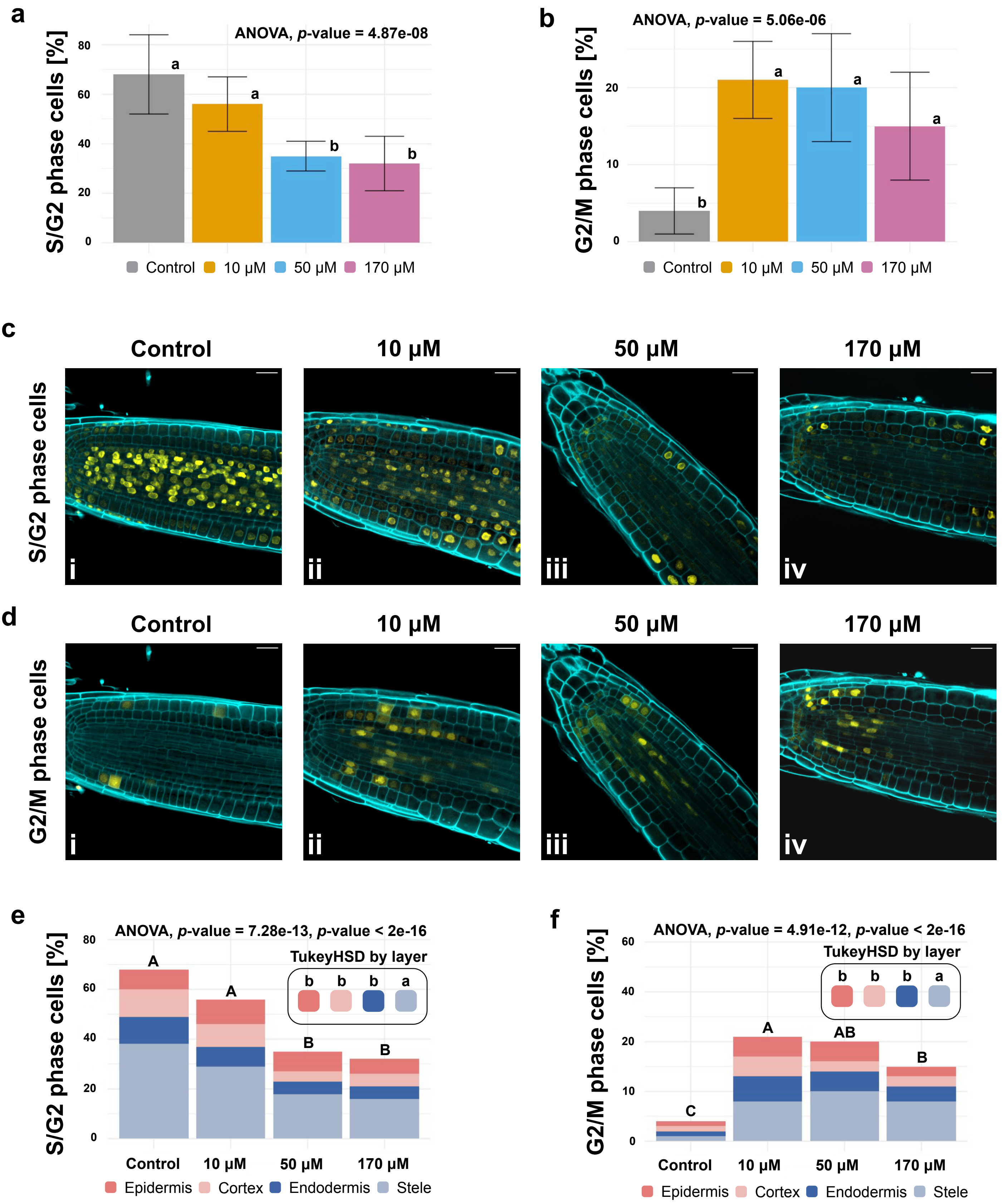
a
ANOVA, p-value = 4.87e-08
S/G2 phase cells [%]
80
60
40
20
0
a
a
b
b
Control
10 µM
50 µM
170 µM
b
ANOVA, p-value = 5.06e-06
G2/M phase cells [%]
20
10
0
b
a
a
a
Control
10 µM
50 µM
170 µM
c
Control
10 µM
50 µM
170 µM
S/G2 phase cells
i
ii
iii
iv
d
Control
10 µM
50 µM
170 µM
G2/M phase cells
i
ii
iii
iv
e
ANOVA, p-value = 7.28e-13, p-value < 2e-16
TukeyHSD by layer
b b b a
S/G2 phase cells [%]
80
60
40
20
0
A
A
B
B
Control
10 µM
50 µM
170 µM
Epidermis
Cortex
Endodermis
Stele
f
ANOVA, p-value = 4.91e-12, p-value < 2e-16
TukeyHSD by layer
b b b a
G2/M phase cells [%]
60
40
20
10
0
C
A
AB
B
Control
10 µM
50 µM
170 µM
Epidermis
Cortex
Endodermis
Stele

**Fig. S8 Evaluation of cell cycle changes in *Arabidopsis thaliana* Cytrap line roots after exposure to 0 µM, 10 µM, 50 µM, 170 µM concentrations of zeocin.** (**a**) Evaluation of S/G2 phase cells relative to total number of cells analyzed in roots, value shown in %. ANOVA revealed statistically significant difference in S/G2 phase cell representation by zeocin concentration ($F(3)=21.73$, $p=4.87e-08$)). Error bars indicate standard deviation. Letters indicate results of TukeyHSD test of with 95% confidence level. (**b**) Evaluation of G2/M phase cells relative to total number of cells analyzed in roots, value shown in %. ANOVA revealed statistically significant difference in G2/M phase cell representation by zeocin concentration ($F(3)=13.7$, $p=5.06e-06$)). Error bars indicate standard deviation. Letters indicate results of TukeyHSD test of two-way ANOVA results with 95% confidence level. (**c**) Representative images showing S/G2 marker expression after root exposure to 0µ M (i), 10µ M (ii), 50µ M (iii), 170µ M (iv) concentrations of zeocin. (**d**) Representative images showing G2/M marker expression after root exposure to 0 µM (i), 10 µM (ii), 50 µM (iii), 170 µM (iv) concentrations of zeocin. (**e**) Evaluation of S/G2 phase cells relative to total number of root cells in selected cell types, value shown in %. Two-way ANOVA revealed statistically significant difference in S/G2 phase cell representation by both zeocin concentration ($F(3)=24.46$, $p=7.28e-13$)) and cell lineage ($F(3)=91.37$, $p<2e-16$)). Letters indicate results of TukeyHSD test of two-way ANOVA results with 95% confidence level. (**f**) Evaluation of G2/M phase cells relative to total number of root cells in selected cell types, value shown in %. Two-way ANOVA revealed statistically significant difference in S/G2 phase cell representation by both zeocin concentration ($F(3)=22.54$, $p=4.91e-12$)) and cell lineage ($F(3)=33.46$, $p<2e-16$)). Letters indicate results of TukeyHSD test of two-way ANOVA results with 95% confidence level. Dataset containing 1002, 816, 800, 998 individual measurements for 0 µM, 10 µM, 50 µM and 170 µM concentrations correspondingly was used for measurements shown on graphs of the figure (**a, b, e, f**).

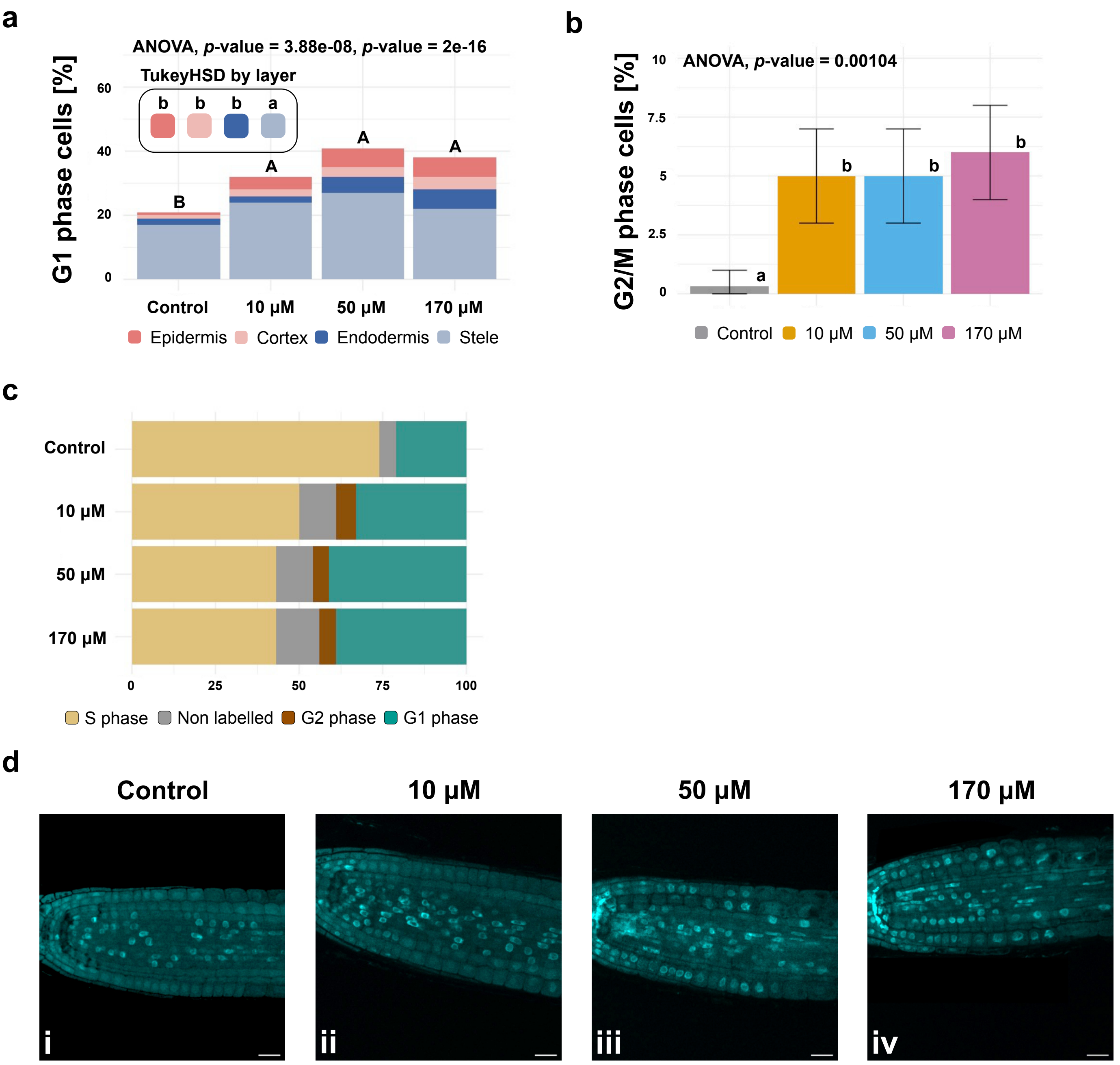

a
ANOVA, p-value = 3.88e-08, p-value = 2e-16
TukeyHSD by layer
b b b a
G1 phase cells [%]
60
40
20
0
B
A
A
A
Control
10 µM
50 µM
170 µM
Epidermis
Cortex
Endodermis
Stele
b
ANOVA, p-value = 0.00104
G2/M phase cells [%]
10
7.5
5
2.5
0
a
b
b
b
Control
10 µM
50 µM
170 µM
c
Control
10 µM
50 µM
170 µM
0
25
50
75
100
S phase
Non labelled
G2 phase
G1 phase
d
Control
10 µM
50 µM
170 µM
i
ii
iii
iv

**Fig. S9 Evaluation of cell cycle changes in *Arabidopsis thaliana* PlaCCI line roots after exposure to 0 µM, 10 µM, 50 µM, 170 µM concentrations of zeocin.** (**a**) Evaluation of G1 phase cells relative to total number of root cells in selected cell types, value shown in %. Two-way ANOVA revealed statistically significant difference in G1 phase cell representation by both zeocin concentration ($F(3)=13.64$, $p=3.88e-08$)) and cell lineage ($F(3)=279.15$, $p<2e-16$)). Letters indicate results of TukeyHSD test of two-way ANOVA results with 95% confidence level. (**b**) Evaluation of G2/M phase cells relative to total number of root cells, value shown in %. ANOVA revealed statistically significant difference in S/G2 phase cell representation by zeocin concentration ($F(3)=10.13$, $p=0.00104$)). Error bars indicate standard deviation. Letters indicate results of TukeyHSD test with 95% confidence level. (**c**) Representation of cells in different phases of the cell cycle, value shown in %. Dataset containing n=2526, n=2438, n=1997, n=1698 cells for 0 µM, 10 µM, 50 µM, 170 µM zeocin respectively was used for measurements shown on graphs of the figure (**a, b, c**). (**d**) Representative images of PlaCCI line showing expression of G1 marker after root exposure to 0 µM (i), 10 µM (ii), 50 µM (iii), 170 µM (iv) concentrations of zeocin. Scale bars, 20 µm.

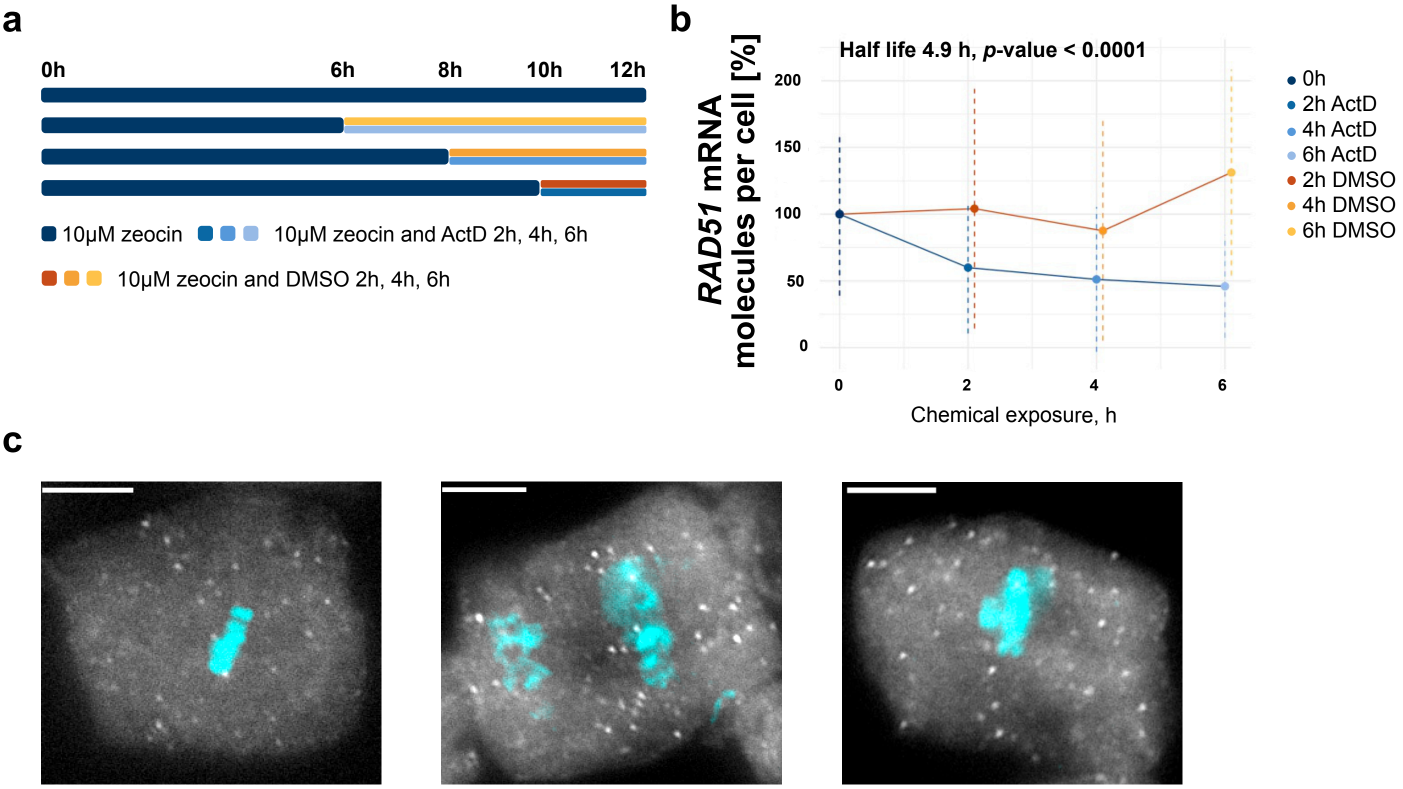
a
0h
6h
8h
10h
12h
10µM zeocin
10µM zeocin and ActD 2h, 4h, 6h
10µM zeocin and DMSO 2h, 4h, 6h
b
Half life 4.9 h, p-value < 0.0001
RAD51 mRNA molecules per cell [%]
200
150
100
50
0
0
2
4
6
Chemical exposure, h
0h
2h ActD
4h ActD
6h ActD
2h DMSO
4h DMSO
6h DMSO
c

**Fig. S10 *RAD51* mRNA molecule half-life evaluation under DNA damage in squashed roots of *Arabidopsis thaliana*.** (**a**) Schematic representation of the experiment. 5-6 days old seedlings were exposed to 10 µM zeocin for 12 hours. A number of seedlings were transferred on media containing 10 µM zeocin and ActD or 10 µM zeocin and DMSO at selected time points. (**b**) *RAD51* mRNA decay evaluation after transcription block with Actinomycin D and in control condition (DMSO). *RAD51* transcription was induced with 10 µM zeocin at all time points. Numbers of mRNA molecules per cell converted to percent, average number of molecules per cell in 12h zeocin exposed sample taken as 100%. Dashed lines indicate standard deviation. Half-life of a transcript shown together with p-value of the function used for calculation. (**c**) Representative images of squashed root cells exposed to 10 µM zeocin showing *RAD51* transcripts in cells undergoing mitosis. Scale bars, 5 µm. Dataset containing n=88, n=79, n=122, n=198, n=131, n=189, n=153 cells for Control, DMSO (2 h, 4 h, 6 h) and ActD (2 h, 4 h, 6 h) respectively.

**Table S1** ***RAD51*** **transcription site (TS) representation in cells residing at G1 and other phases of the cell cycle in *Arabidopsis thaliana* CDT1-CFP line roots.**

| Sample | Total number of cells analyzed | Cells with TS signals, % | G1 cells with TS signals, % | non G1 cells with TS signals, % |
|---|---|---|---|---|
| **Control** | 1096 | 1% | 0,5% | 0,5% |
| **10 µM** | 1088 | 3,4% | 1,7% | 1,7% |
| **50 µM** | 1054 | 12,2% | 6,8% | 5,4% |
| **170 µM** | 422 | 15,2% | 7,1% | 8,1% |
